# Epigenetic Regulation of Chondrogenesis: JMJD3 and UTX as Key Targets for Gene-Modified Mesenchymal Stem Cell Therapy in Cartilage Tissue Engineering

**DOI:** 10.1101/2025.06.06.658259

**Authors:** L Allas, J Aury-Landas, Q Rochoux, A Julien, E Lhuissier, M Lente, S Brochard, A Veyssiere, K Boumédiene, C Baugé

**Author notes:** Université de Caen Normandie, CNRS, Normandie Université, ISTCT UMR6030, GIP CYCERON, F-14000 Caen, France. Address for correspondence: Catherine BAUGE UR7451 BioConnecT Université de Caen Normandie 14032 CAEN FRANCE.

## Abstract

Osteoarthritis is a major cause of disability in older adults, and among the promising therapeutic strategies, cartilage tissue engineering shows great potential. Histone methylation plays a critical role in cartilage development, making it an appealing target for improving cartilage engineering protocols. In this study, we evaluated the roles of JMJD3 (KDM6B) and UTX (KDM6A), both demethylases of histone H3 at lysine 27 (H3K27), in chondrogenesis and their application in gene-modified mesenchymal stem cell therapy for cartilage tissue engineering.

Using high-throughput analyses such as ChIP-Seq and whole-transcriptome microarray, we explored the functions of JMJD3 and UTX in human bone marrow-derived mesenchymal stem cells (hBM-MSC) undergoing chondrogenesis. We investigated the impact of inhibiting JMJD3 and UTX with the pharmacological inhibitor GSK-J4 or using siRNA. Additionally, the effects of transiently transfecting JMJD3 or UTX expression vectors were assessed both *in vitro* and *in vivo*, following the implantation of hBM-MSC embedded in alginate in nude mice.

Our findings revealed that JMJD3 is specifically upregulated during chondrogenesis in hBM-MSC, and is crucial for this differentiation process. In contrast, UTX was found to be dispensable for chondrogenesis. Nevertheless, both JMJD3 and UTX share the ability to demethylate similar gene loci, thereby promoting the expression of chondrogenic signature genes, which suggests functional redundancy. Notably, the genes encoding these H3K27me3 demethylases emerge as strong candidates for enhancing gene-modified mesenchymal stem cell therapy for cartilage tissue engineering, as their overexpression during chondrogenesis significantly increased the formation of thicker cartilage discs enriched with type II collagen.

In conclusion, this study provides important insights into the epigenetic regulation of chondrogenesis, especially regarding the role of H3K27me3 demethylases. We demonstrate that, although JMJD3 and UTX have overlapping targets, only JMJD3 is critical for the chondrogenesis process. Additionally, the findings emphasize the potential of transient JMJD3 transduction, along with a lesser emphasis on UTX, as effective strategies for improving gene-modified mesenchymal stem cell therapy in cartilage tissue engineering.

## Introduction

Osteoarthritis is a leading cause of disability worldwide, profoundly affecting the quality of life for millions of individuals. Current therapeutic strategies for this debilitating rheumatic disease are primarily symptomatic, focusing on pain relief and improved joint mobility, without achieving cartilage regeneration. The limited self-repair capacity of cartilage, due to its avascular nature, underscores the urgent need for innovative regenerative approaches. Consequently, tissue engineering strategies have garnered significant attention as promising solutions for cartilage repair and regeneration.

The introduction of autologous chondrocyte transplantation by Mat Brittberg in 1994 marked a pivotal advancement in cell-based cartilage repair^1^. This technique involves harvesting chondrocytes from articular cartilage, expanding them *in vitro*, and re-implanting them into cartilage defects. However, the limited yield of chondrocytes from donor tissue necessitates extensive *in vitro* expansion, during which chondrocytes tend to lose their native phenotype and acquire a fibroblast-like morphology^2^. These phenotypic alterations compromise the biomechanical properties of the regenerated cartilage and, consequently, its clinical efficacy.

To overcome these limitations, embryonic and adult stem cells, particularly mesenchymal stem cells (MSCs), have emerged as attractive alternatives. MSCs exhibit robust chondrogenic potential, self-renewal capabilities, and immunomodulatory properties^3,4^, making them ideal candidates for cartilage tissue engineering. Despite these advantages, achieving and maintaining a stable chondrocyte phenotype post-differentiation remains a significant challenge.

Epigenetic mechanisms, including DNA methylation, histone modifications (e.g., methylation and acetylation), and noncoding RNAs, are critical regulators of stem cell maintenance and lineage commitment. These mechanisms are dynamic and reversible, positioning them as compelling targets for therapeutic interventions^3^. Understanding the epigenetic signatures associated with MSC differentiation is crucial for developing strategies to promote efficient chondrogenesis and facilitate cartilage regeneration using cell-based therapies.

Among various epigenetic regulators, histone modifications play a central role in modulating chromatin dynamics and gene expression. Specifically, histone H3 lysine 27 (H3K27) methylation has been primarily associated with transcriptional repression. This modification is catalyzed by the histone methyltransferase EZH2 and can be reversed by two lysine-specific demethylases (KDMs): Ubiquitously Transcribed Tetratricopeptide Repeat, X chromosome (UTX, also known as KDM6A) and JmjC Domain-Containing Protein 3 (JMJD3, also referred to as KDM6B).

While the role of EZH2 in chondrogenesis and chondrocyte function has been extensively explored, the specific contributions of UTX and JMJD3 remain less defined. JMJD3 is highly expressed in chondrocyte lineage cells during endochondral ossification, with its deletion leading to delayed skeletal development in mice^5^. In contrast, UTX deletion has been associated with increased articular cartilage thickness and attenuated osteoarthritic symptoms^6^, suggesting distinct and potentially opposing roles for these enzymes in cartilage biology. Paradoxically, some studies indicate that both JMJD3 and UTX may exacerbate cartilage degeneration by impairing chondrocyte anabolism^6–8^, highlighting the complexity of their functions.

In this study, we investigated the regulation and functional roles of UTX and JMJD3 during *in vitro* chondrogenesis of human bone marrow-derived mesenchymal stem cells (hBM-MSCs). Through gain- and loss-of-function approaches, we analyzed the specific contributions of these H3K27me3 demethylases to the chondrogenic differentiation process. Additionally, we employed chromatin immunoprecipitation sequencing (ChIP-Seq) to identify their genome-wide binding targets and assessed their potential therapeutic applications in cartilage tissue engineering *in vivo*.

## Material and methods

### Stem cells isolation and amplification

Bone marrow was obtained from the iliac crests of adult female donors (ages 53–77 years; median age 64 years) who underwent hip replacement surgery. All patients provided written informed consent, and the study was approved by the local ethics committee named “Comité de Protection des Personnes Nord Ouest III” (agreement #A13-D46-VOL.19). Cells were isolated as previously described^9^. The bone marrow was fractionated on a Hypaque-Ficoll (GE Life Sciences, Velizy-Villacoublay, France) density gradient to isolate mononuclear cells. These cells were seeded at a density of 5 × 10⁴ cells/cm² and cultured in alpha-MEM supplemented with 10% fetal bovine serum (Lonza, Levallois-Perret, France), 2 mM L-glutamine, 1 ng/ml FGF-2 (Merck, Loiret, France), and antibiotics (penicillin-streptomycin, Lonza) at 37°C in 5% CO₂.

At confluency, cells were harvested by trypsinization (0.25% trypsin/1 mM EDTA, Invitrogen) and re-seeded at 10³ cells/cm². Cells were used at passages 3 to 5. Following amplification, the differentiation capacity and colony-forming unit potential of hBM-MSCs were confirmed. Additionally, the absence of mycoplasma DNA contamination was verified by PCR.

### Chondrogenesis

Chondrogenesis was induced by culturing cells in a 3D alginate matrix using chondrogenic medium, as previously described^9^. Briefly, MSCs were suspended in alginate (Kelco) at a density of 5 × 10⁶ cells/ml and dropped into a CaCl₂ solution to form beads. The beads were then cultured in chondrogenic medium containing DMEM supplemented with antibiotics (penicillin-streptomycin, Lonza), 100 nM dexamethasone (Merck), 50 µg/ml ascorbic acid-2-phosphate (Merck), 40 µg/ml proline (Merck), 1 mM sodium pyruvate (Merck), a 1:100 dilution of Insulin-Transferrin-Selenium (ITS) +1 supplement (Merck), and 10 ng/ml TGF-β3 (Merck). After the culture period, the alginate beads were dissolved in a solution of 55 mM sodium citrate and 150 mM NaCl, followed by gentle centrifugation to isolate cells for further analysis.

### ChIP-Seq

The ChIP-Seq experiment has been conducted from chromatin of two different patients by Diagenode ChIP-seq/ChIP-qPCR Profiling service (Diagenode Cat# G02010000). The chromatin was prepared using the iDeal ChIP-seq kit for Histones (Diagenode Cat# C01010059). Chromatin was sheared using Bioruptor® Pico sonication device (Diagenode Cat# B01080010) combined with the Bioruptor® Water cooler for 8 cycles using a 30’’ [ON] 30’’ [OFF] settings. Shearing was performed in 0.65 ml Bioruptor® Pico Microtubes (Diagenode Cat# C30010011) with the following cell number: 400 000 in 100μl. 20μl of this chromatin was used to assess the size of the DNA fragments obtained by High Sensitivity NGS Fragment Analysis Kit (DNF-474) on a Fragment Analyzer™ (Advanced Analytical Technologies, Inc.). ChIP was performed using IP-Star® Compact Automated System (Diagenode Cat# B03000002) following the protocol of the aforementioned kit. Chromatin corresponding to approximately 250 000 cells was immunoprecipitated with H3K4me3 or H3K27me3 antibodies. qPCR analyses were made to check ChIP efficiency using KAPA SYBR® FAST (Sigma-Aldrich) on LightCycler® 96 System (Roche) and results were expressed as % recovery = 2(Ct_input-Ct_sample) / 100*2((Ct_input-6.64)-Ct_sample) / 100*2((Ct_input-3.32)-Ct_sample). Primers used were the following:GAPDH-TSS (Diagenode Cat# C17011047-50) and TSH2B (Diagenode Cat# C17011041-50).

Libraries were prepared using IP-Star® Compact Automated System (Diagenode Cat# B03000002) from input and ChIP’s DNA using MicroPlex Library Preparation Kit v2 (12 indices) (Diagenode Cat# C05010013). Library amplification is assessed using High Sensitivity NGS Fragment Analysis Kit (DNF-474) on a Fragment Analyzer™ (Advanced Analytical Technologies, Inc.). Libraries were then purified (double size-selected) using Agencourt® AMPure® XP (Beckman Coulter) and quantified using Qubit™ dsDNA HS Assay Kit (Thermo Fisher Scientific, Q32854). Finally, the fragment size was analyzed by High Sensitivity NGS Fragment Analysis Kit (DNF-474) on a Fragment Analyzer™ (Advanced Analytical Technologies, Inc.).

Quality control of sequencing reads was performed using FastQC. Reads were aligned to the reference genome (hg38) obtained from the UCSC genome browser using BWA. Samples were filtered for regions blacklisted by the ENCODE project. Multimapping reads and PCR duplicates were removed using samtools functionalities. Alignment coordinates were converted to BED format using BEDTools v.2.17 and peak calling was performed using MACS2. Peaks were visualized on genome using Integrated Genome Viewer (IGV). A simple binary comparisons (identifying common or unique peaks) were performed; and functional enrichments in Annoted Keywords (UniProt) from genes having unique peaks for considered condition were performed and visualized using STRING database (version 12.0; ^10^_)._

### RNA isolation and real-time reverse transcription-polymerase chain reaction (RT-PCR)

RNA was extracted using the NucleoSpin® RNA kit (Macherey-Nagel, Hoerdt, France) following the manufacturer’s protocol. After extraction, the RNA was treated with DNase and reverse-transcribed into cDNA, as previously described^11^. Briefly, reverse transcription was performed in a reaction mixture containing 1 µg of RNA, oligo dT primers (2.5 µM), 500 µM dNTPs, 1× First Strand Buffer, and 10 U/μL Moloney Murine Leukemia Virus reverse transcriptase (M-MLV-RT, Invitrogen, Courtaboeuf, France). The reaction was conducted at 37°C for 50 minutes, followed by enzyme inactivation at 70°C for 15 minutes, using a T100™ Thermal Cycler (Bio-Rad, Les Ulis, France). The resulting cDNA was amplified by real-time PCR using the StepOnePlus™ Real-Time PCR System (Applied Biosystems, Courtaboeuf, France) with the following primers: COL2A1-F: ACTGGATTGACCCCAACCAA; COL2A1-R: TTGGGAACGTTTGCTGGATT; RPL13A-F: GAGGTATGCTGCCCCACAAA; RPL13A-R: GTGGGATGCCGTCAAACAC; ACAN-F: TCGAGGACAGCGAGGCC; ACAN-R: TCGAGGGTGTAGCGTGTAGAGA SOX9-F: CCCATGTGGAAGGCAGATG; SOX9-R TTCTGAGAGGCACAGGTGACA; SOX5-F: ATCCCAACTACCATGGCAGCT; SOX5-R: TGCAGTTGGAGTGGGCCTA; SOX6-F: GCAGTGATGAACATGTGGCCT; SOX6-R: CGTGTCCCAGTCAGCAGCATCT; UTX-F: TACAAATCCCGAACAACCC; UTX-R: TGAGGAGGCCTGGTACTGT; JMJD3-F: AGCTGGCCCTGGAACGATA; JMJD3-R: GGCCCTGGTAAGCATTT. The relative mRNA level was calculated with the 2^−ΔΔCT^ method. RPL13a was used as reference.

### Gene expression microarray analysis

RNA integrity was assessed using a 2100 Bioanalyzer (Agilent Technologies, Les Ulis, France) and RNA 6000 Nano chips (Agilent Technologies), following the manufacturer’s instructions. RNA integrity numbers (RINs) above 9 were considered suitable for microarray analysis. Two-color microarray-based gene expression analysis was performed according to the manufacturer’s protocol (Agilent Technologies). Briefly, 100 ng of total RNA was amplified and labeled using the Low Input Quick Amp Labeling Kit, Two-Color (Agilent Technologies), and hybridized to a Human Gene Expression 8 × 60K v3 Microarray (design ID 072363, Agilent Technologies). Slides were scanned using a G2505C Microarray Scanner (Agilent Technologies). Raw data were extracted and Lowess-normalized using Feature Extraction Software (v. 10.7.3, Agilent Technologies) and further analyzed with GeneSpring GX software (v. 13.1.1, Agilent Technologies). Microarray probes with signals that were either not positive and significant or below the background were filtered out. Genes with a fold changes > 1.5 were considered differentially expressed genes (DEGs). Functional enrichment from DEG was visualized using STRING databases (v. 12.0)^10^. A False discovery rate (FDR) < 0.05 were considered as significant. Only the 10 first terms were shown (category: “KEGG pathways” or “Biological Process”; merge: “Merge rows by term similarity ≥ 0.8”).

### Plasmids, siRNAs and transfection

To overexpress UTX and JMJD3, pCMV-HA-UTX (#24168, Addgene) and pCMV-HA-JMJD3 (#24167, Addgene) were used, respectively^12^. Pmax-GFP was supplied by Amaxa. Pools of four control siRNAs (D-001810-10-50) or siRNAs targeting JMJD3 (L-023013-01-0005) or UTX (L-014140-01-0005) mRNA were obtained from Dharmacon.

For transfection, MSCs were harvested and resuspended in P1 Primary Cell 4D-Nucleofector™ X Solution (Amaxa), supplemented with either plasmid DNA or siRNA. Mixtures of 4 × 10⁶ cells with 8 µg of DNA or 4 × 10⁶ cells with 400 pmol of siRNA were placed in 1 cm² transfection cuvettes and nucleoporated using the FF-104 predefined program on a 4D-Nucleofector™ X Unit (Amaxa). Ten minutes after electroporation, cells were encapsulated in alginate beads, transferred to six-well plates, and cultured in chondrogenic medium.

### Protein extraction and western blotting

Cells were rinsed and then scraped into Radioimmunoprecipitation Assay (RIPA) buffer supplemented with phosphatase and protease inhibitors. The protein extracts (30 μg) were subjected to fractionation using 10% sodium dodecyl sulfate–polyacrylamide gel electrophoresis (SDS-PAGE), followed by transfer to polyvinylidene difluoride (PVDF) membranes (Bio-Rad, Marnes-la-Coquette, France). Western blotting was performed as previously described^13^. The following antibodies were used: JMJD3 (#3457) and UTX (D3Q1I) from Cell Signaling, β-actin (sc-47778), goat anti-mouse IgG-HRP (sc-2005), and goat anti-rabbit IgG-HRP (sc-516087) from Santa Cruz Biotechnology.

### *In vivo* experiments

Animal experimental procedures were conducted in accordance with local legislation and approved by the ethics committee (Comité d’Ethique Normandie en Matière d’Expérimentation Animale, agreement #19573), with authorization from the French Ministry of Research.

A total of 10 NMRI-Foxn1 nu/nu mice (7 weeks old, male, provided by Janvier Labs) were housed at 5 per cage under a 12-hour light/dark cycle, with ad libitum access to water and food. The animals were kept in a germ-free facility (ONCOModels, 30 m², Cyceron, Caen, France). Throughout the experiment, each animal was handled humanely in accordance with internationally accepted ethical guidelines for laboratory animal use and care, and all efforts were made to minimize animal suffering.

Cellularized alginate beads were implanted subcutaneously in nude mice under sterile conditions. During the procedure, mice were anesthetized with 2% isoflurane. A 5 mm incision was made in the dorsal region, followed by a second incision in the lumbar region of the skin. Two subcutaneous pouches were created on each side of the lesions to accommodate the beads. To reduce animal use, each mouse received four beads: one with untransfected cells (Ctrl), one with JMJD3-transfected cells, one with UTX-transfected cells, and one with GFP-transfected cells.

After 4 weeks, the mice were anesthetized and euthanized using CO_2_. Surface of beads were evaluated using the formula: A=π×(length/2)×(width/2). Tissue specimens were collected and iced for collagen assay, or embedded in Optimal Cutting Temperature (OCT) for immunohistofluorescence.

### Insoluble collagen assay

Collagen content was quantified using the Sircol Insoluble Collagen Assay Kit (S2000, Biocolor) following the manufacturer’s protocol. Absorbance was measured at 550 nm using a Multiskan GO spectrophotometer (Thermo Scientific).

### Immunohistofluorescence

Sections with a thickness of 10 µm were prepared using a cryostat (CM3050 S, Leica). The slides were thawed and incubated at room temperature for 30 minutes. For immunofluorescence, sections were washed once with Phosphate Buffered Saline (PBS, Lonza) for 5 minutes. The slides were then incubated overnight at 4°C in a solution containing PBS, 0.25% Triton X-100, 1% BSA, and primary antibodies diluted as follows: Collagen II (1:500, ab34712, Abcam), or Aggrecan (1:500, AB1031, Merck Millipore). After incubation, the slides were washed and incubated for 1 hour at room temperature in a secondary antibody solution containing PBS, 0.25% Triton X-100, 1% BSA, Hoechst solution (1:1000), and Alexa Fluor 594 Goat IgG Anti-Rabbit IgG (1:400, 111-585-003, Jackson ImmunoResearch). Fluorescence was evaluated using the EVOS FL Auto 2 Cell Imaging System (ThermoFisher Scientific, Illkirch-Graffenstaden, France).

### Statistical analysis

Statistical analyses were carried out on the GraphPad Prism 8 software. After checking the normal distribution of samples, t-test or one-way ANOVA tests were used for simple or multiple comparisons. Results are expressed as the mean value of three or four patients (biological replicates) ± the standard error of the mean (SEM). The ROUT method was used to determine outliers (Q = 1%). For all analysis, p-values < 0.05 were considered significant.

## Results

### H3K27 methylation was decreased genome-wide during chondrogenesis

To induce chondrogenesis, human bone marrow-derived mesenchymal stem cells were encapsulated in alginate beads and cultured in a differentiation medium supplemented with TGF-β3 for seven days. Chromatin immunoprecipitation followed by deep sequencing (ChIP-Seq) was employed to analyze genome-wide enrichment of the permissive histone mark H3K4me3 and the repressive mark H3K27me3 (Figure 1A).

**Figure 1:**
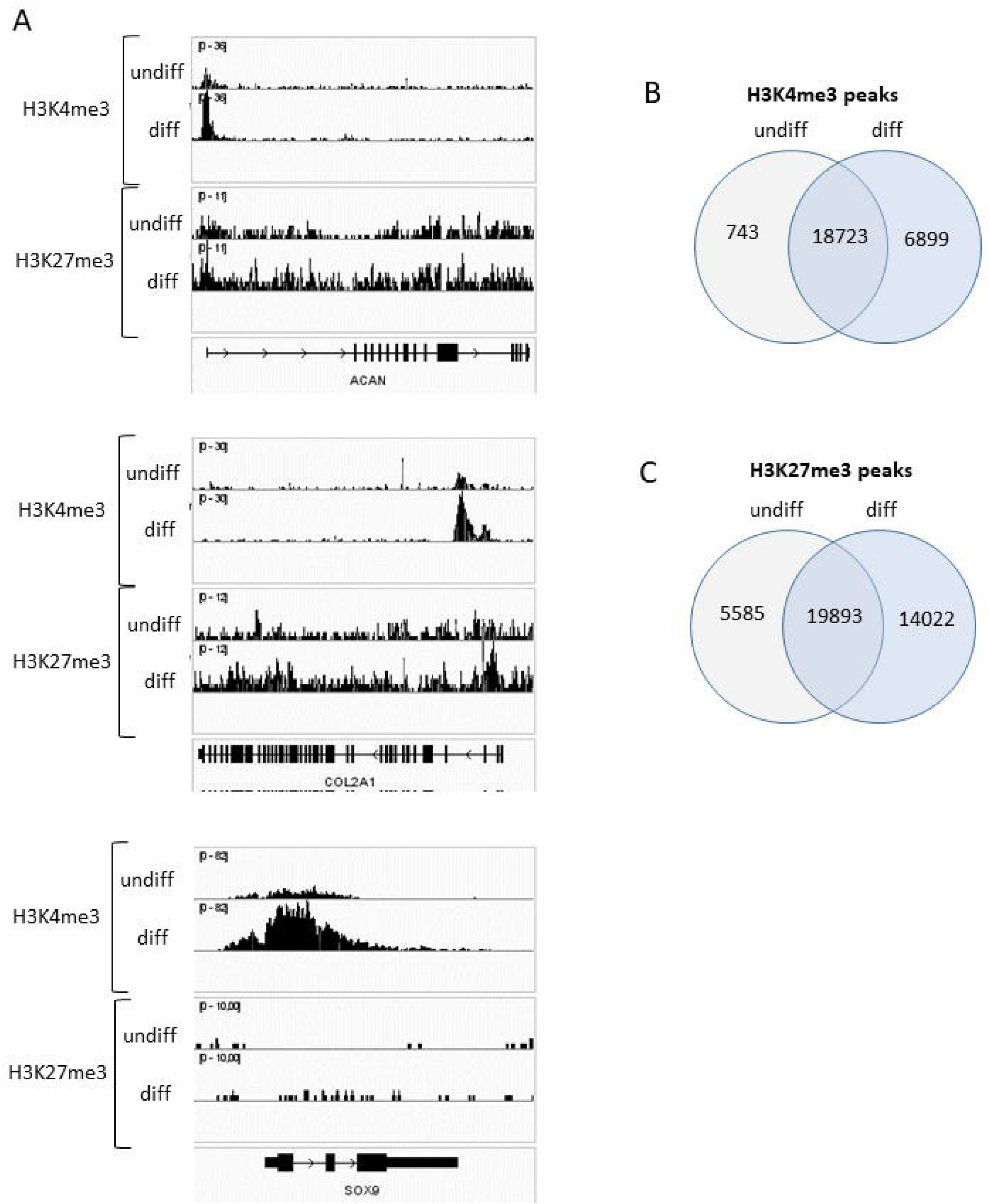
H3K27 methylation is locally decreased during chondrogenesis. Human bone marrow-derived mesenchymal stem cells (hBM-MSCs) were embedded in alginate beads and cultured in a differentiation medium containing TGF-β3 for 7 days. Chromatin immunoprecipitation followed by sequencing (ChIP-Seq) was performed to analyze genome-wide H3K4me3 and H3K27me3 modifications. (A) Representative loci of chondrogenic signature genes (ACAN, COL2A1, and SOX9) are shown. (B) Venn diagram illustrating the number of common and unique H3K4me3 peaks between differentiated and undifferentiated conditions. (C) Venn diagram showing the common and unique H3K27me3 peaks between the two conditions.

In undifferentiated MSCs, H3K27me3 was broadly distributed across the COL2A1 locus, while H3K4me3 was confined to a peak at the first introns, indicative of a bivalent or inactive gene state. Upon chondrogenic differentiation (day 7), H3K4me3 enrichment increased, corresponding to the activation of COL2A1 transcription. However, H3K27me3 enrichment persisted, suggesting incomplete resolution of the repressive mark. Similarly, in undifferentiated MSCs, H3K4me3 peaks were detected at the ACAN promoter and across a broader region of SOX9, spanning from upstream of the promoter to the second intron. These peaks intensified following chondrogenesis, consistent with the upregulation of these genes. Despite these transcriptional changes, H3K27me3 remained enriched across the ACAN locus in both conditions, indicating that differentiation was not fully complete across all cells.

Genome-wide occupancy analysis of ChIP-Seq data identified 26 365 H3K4me3 sites and 39 500 H3K27me3 sites (Figures 1B and 1C). A substantial proportion of these sites (18 723 for H3K4me3 and 19 893 for H3K27me3) were common to both undifferentiated and differentiated states (*i.e.* present in all replicates of undifferentiated MSCs and all replicates of differentiated cells). However, we identified 743 H3K4me3 sites (2.8%) and 5 585 H3K27me3 sites (14.1%) unique to undifferentiated MSCs, and thus not present in differentiated cells. This indicates that over 10% of H3K27me3 sites were demethylated during chondrogenesis, underscoring the pivotal role of H3K27me3 demethylation in this differentiation process.

Collectively, these findings indicate that although locus-specific H3K27me3 changes at COL2A1 or SOX9 were not detected at the examined time point, a genome-wide reduction in H3K27me3 occurred during chondrogenic differentiation. This suggests the critical involvement of H3K27me3 demethylases in mediating chondrogenic commitment.

### The pharmacological inhibition of histone demethylases JMJD3 and UTX impaired chondrogenesis by altering H3K27me3 demethylation

To elucidate the role of JMJD3 and UTX in modulating the H3K27me3 landscape during chondrogenic differentiation, we employed GSK-J4, a potent inhibitor of JMJD3 and UTX^14^ (Figure 2A). GSK-J4 acts by competing with essential cofactors, 2-oxoglutarate (also known as α-ketoglutarate) and Fe^2+^, which are critical for the catalytic activity of these demethylases.

**Figure 2:**
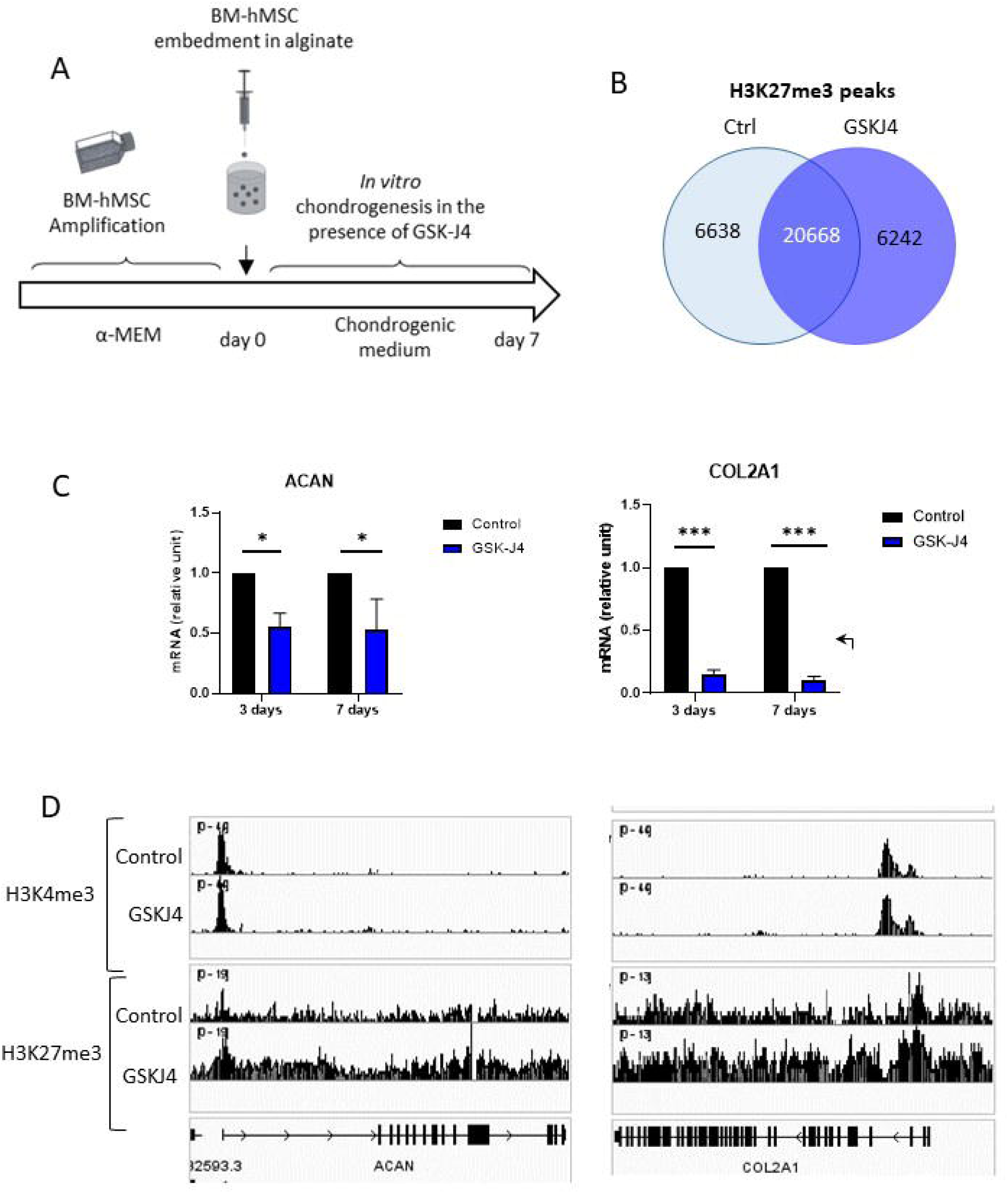
GSK-J4 alters H3K27me3 demethylation during chondrogenesis and impairs COL2A1 and ACAN expression. hBM-MSCs were cultured in alginate beads in chondrogenic medium with or without GSK-J4 for 7 days. (A) Design of the experiment. (B) Chromatin immunoprecipitation followed by sequencing (ChIP-seq) was performed to analyze genome-wide H3K4me3 and H3K27me3 modifications. Venn diagram illustrating the number of common and unique H3K27me3 peaks between the conditions. (C) RNA was extracted, and the mRNA expression levels of COL2A1 and ACAN were quantified by real-time RT-PCR. Data are expressed as mean ± SEM (n=3). *: pvalue < 0.05; ***: pvalue < 0.001. ( D) H3K4me3 and H3K27me3 peaks at the loci of these genes are shown.

Treatment with GSK-J4 markedly altered the H3K27me3 profile in differentiating cells (Figure 2B). Approximately 20% of H3K27me3-enriched sites were consistently detected across all GSK-J4-treated replicates but were absent in the control group, suggesting that GSK-J4 either promotes methylation at these sites or, more likely, impairs their demethylation. This finding indicates that pharmacological inhibition of H3K27me3 demethylases can influence chondrogenesis.

To validate this hypothesis, we evaluated the expression of key chondrogenic marker genes using RT-PCR. GSK-J4 treatment significantly inhibited the expression of ACAN and COL2A1 (Figure 2C), correlating with increased H3K27me3 enrichment at their loci (Figure 2D). In contrast, H3K4me3 levels at these genes remained unaffected by GSK-J4, highlighting the specificity of GSK-J4’s effect on H3K27me3.

Additionally, whole-transcriptome analysis was conducted to comprehensively assess the impact of GSK-J4 during hBM-MSC chondrogenesis. We identified 396 differentially expressed genes (DEGs) with a fold change greater than 1.5 (Table 1), of which 254 (64%) were downregulated. This pattern aligns with a global increase in the repressive H3K27me3 mark following JMJD3/UTX inhibition. Functional enrichment analysis of DEGs revealed significant associations with biological processes and pathways related to “chondrocyte differentiation”, “skeletal system development,” “regulation of Wnt signaling pathway,” and “TGF-beta signaling pathway” (Figures 3A and 3B). These enrichments predominantly include downregulated genes, reinforcing the critical role of H3K27me3 demethylation in chondrogenesis. Pathways associated with “response to copper ion”? “detoxification of copper ion” and “Mineral absorption” were enriched, primarily involving genes upregulated by GSK-J4 during chondrogenesis (Suppl tables 1 and 2).

**Figure 3:**
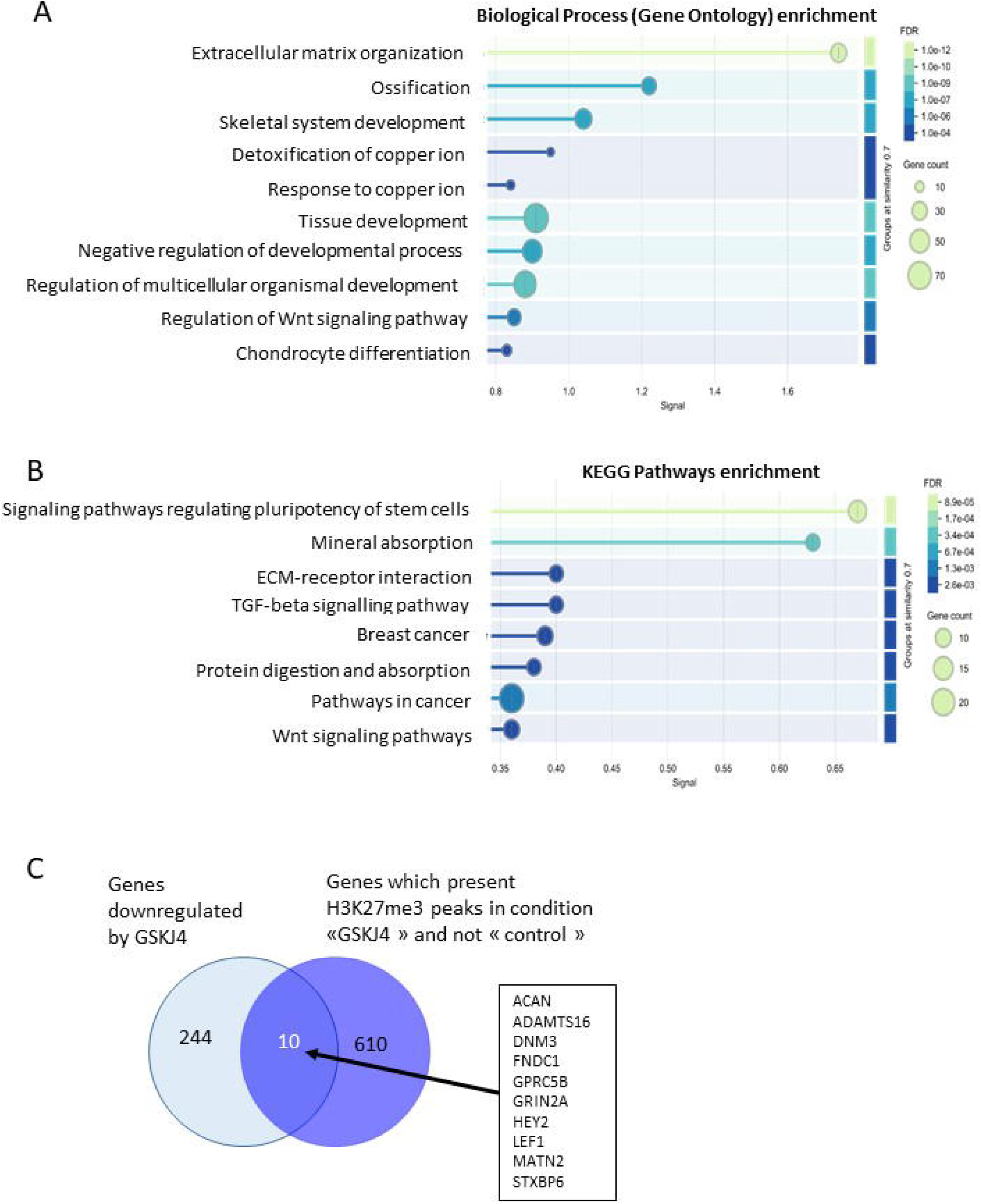
GSK-J4 regulates genes Involved in the extracellular matrix and cartilage formation. hBM-MSCs were cultured in alginate beads in chondrogenic medium with or without GSK-J4 for 7 days. Total RNA was extracted, and transcriptome analysis was performed using RNA microarrays to identify genes whose expression was deregulated by GSK-J4. (A, B) Gene Ontology (GO) biological process enrichment and KEGG pathway analyses were conducted and visualized using STRING tools. (C) Venn diagram showing the overlap of genes that were downregulated and exhibited increased H3K27me3 methylation in the presence of GSK-J4 during chondrogenesis.

Next, to explore the relationship between DEGs and H3K27me3 changes, we integrated transcriptomic data with ChIP-Seq results. As expected, several downregulated genes exhibited new H3K27me3 peaks at their promoters following GSK-J4 treatment (Figure 3C). These genes included ACAN, HEY2, and LEF1, which are key regulators of chondrogenesis.

Collectively, these transcriptomic and ChIP-Seq analyses comparing hBM-MSCs undergoing chondrogenesis with or without GSK-J4 treatment suggest that UTX and/or JMJD3 demethylate H3K27me3 at loci of chondrogenesis-related genes. This demethylation facilitates gene derepression, thereby promoting chondrogenic differentiation.

### JMJD3, but not UTX, was upregulated during chondrogenesis and was required for mesenchymal stem cell differentiation into chondrocytes

Furthermore, we investigated whether the effects of GSK-J4 were due to the specific inhibition of JMJD3 and/or UTX. We analyzed the expression of these demethylases during bone marrow-derived mesenchymal stem cell differentiation into chondrocytes. We observed that only JMJD3 expression was significantly upregulated during chondrogenesis, while UTX expression remained unchanged (Figure 4A). This differential expression pattern suggests a potential role for JMJD3, but not UTX, in chondrogenic differentiation.

**Figure 4:**
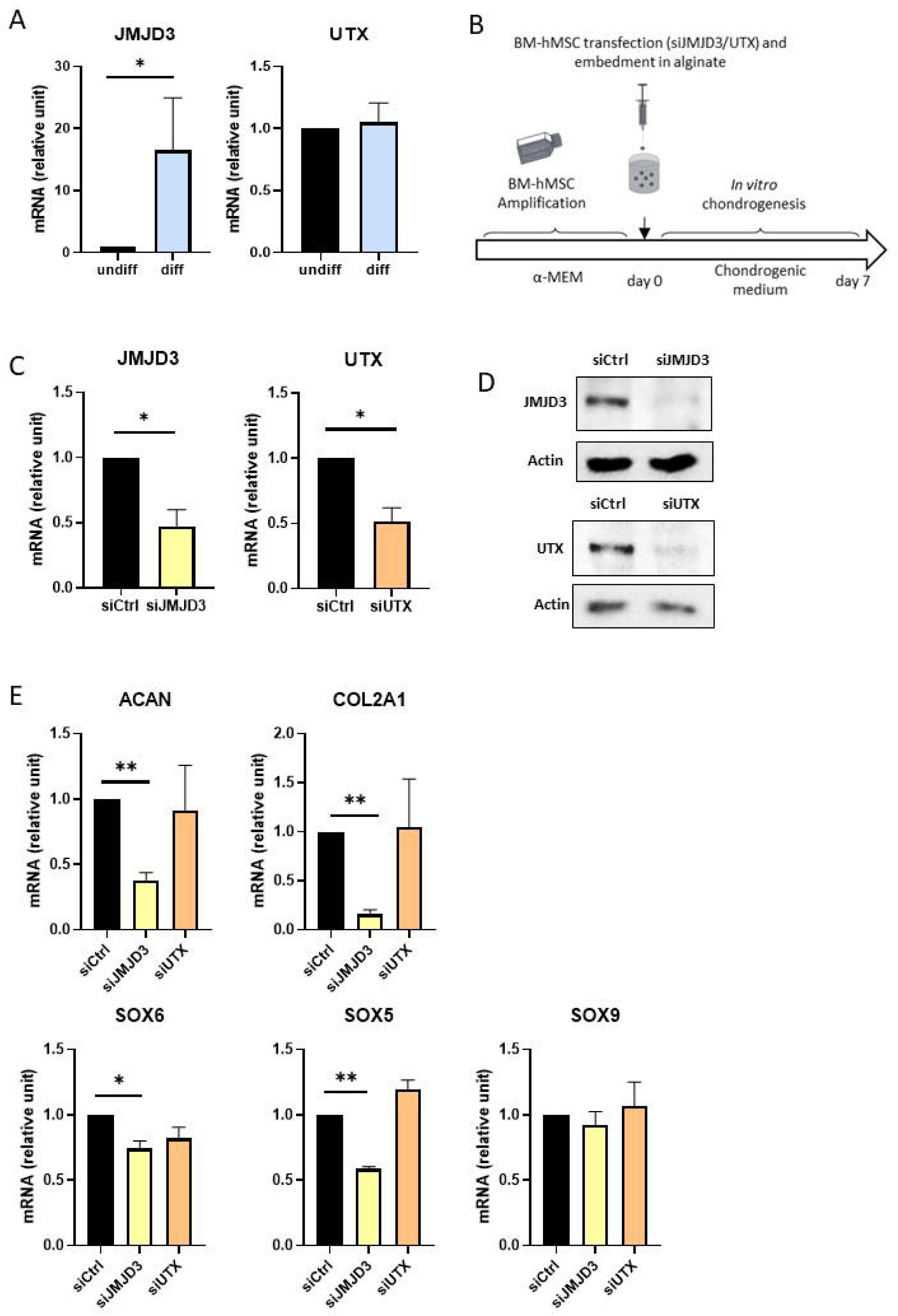
JMJD3, but not UTX, is upregulated during chondrogenesis and required for mesenchymal stem cell differentiation into chondrocytes. (A) hBM-MSCs were embedded in alginate beads and cultured in chondrogenic differentiation medium (diff) or α-MEM (undiff) for 7 days. Total RNA was extracted, and the expression of JMJD3 and UTX was evaluated by RT-PCR (*n* = 8). (B) For knockdown experiments, hBM-MSCs were nucleofected with control siRNA (siCtrl), siUTX, or siJMJD3 and cultured in chondrogenic medium for 7 days. (C, D) Protein and RNA were extracted to assess the expression of JMJD3 and UTX at both mRNA (C) and protein levels (D) using RT-PCR and western blotting, respectively. (E) mRNA expression levels of SOX5, SOX6, SOX9, ACAN, and COL2A1 were quantified by RT-PCR (*n* = 3). Data are expressed as mean ± SEM. *: pvalue < 0.05; **: pvalue < 0.01.

To test this hypothesis, we performed targeted knockdown of JMJD3 and UTX using small interfering RNA (siRNA) (Figure 4B). The siRNA-mediated knockdown effectively reduced JMJD3 and UTX expression at both mRNA and protein levels during chondrogenesis (Figures 4C and 4D). However, subsequent analysis of key chondrogenic marker genes revealed that only JMJD3 knockdown significantly decreased the expression of SOX5, SOX6, ACAN, and COL2A1 (Figure 4E). In contrast, UTX knockdown had no appreciable effect on the expression of these genes. These findings demonstrate that JMJD3, but not UTX, is essential for the chondrogenic differentiation of hBM-MSCs, highlighting pivotal role of JMJD3 in regulating the transcriptional program required for chondrocyte development.

### Both JMJD3 and UTX ectopic expression enhanced in vitro chondrogenesis

Given that JMJD3 knockdown impaired chondrogenesis, we hypothesized that its overexpression would have the opposite effect, promoting chondrogenic differentiation. To test this, we transiently transfected hBM-MSCs with expression vectors for JMJD3 or GFP (as a control) prior to inducing chondrogenesis for 7 days. Our results supported our hypothesis, revealing that JMJD3 overexpression significantly enhanced the mRNA expression of key chondrogenic markers ACAN and COL2A1 (Figure 5A). Interestingly, we also evaluated the effect of UTX overexpression. Despite UTX not being essential for chondrogenesis based on knockdown experiments, its forced expression similarly to JMJD3 led to an increase in ACAN and COL2A1 expression. These findings suggest that while only JMJD3 is indispensable for chondrogenic differentiation, both JMJD3 and UTX can enhance chondrogenesis when ectopically expressed, indicating their potential complementary roles in promoting chondrogenic gene expression.

**Figure 5:**
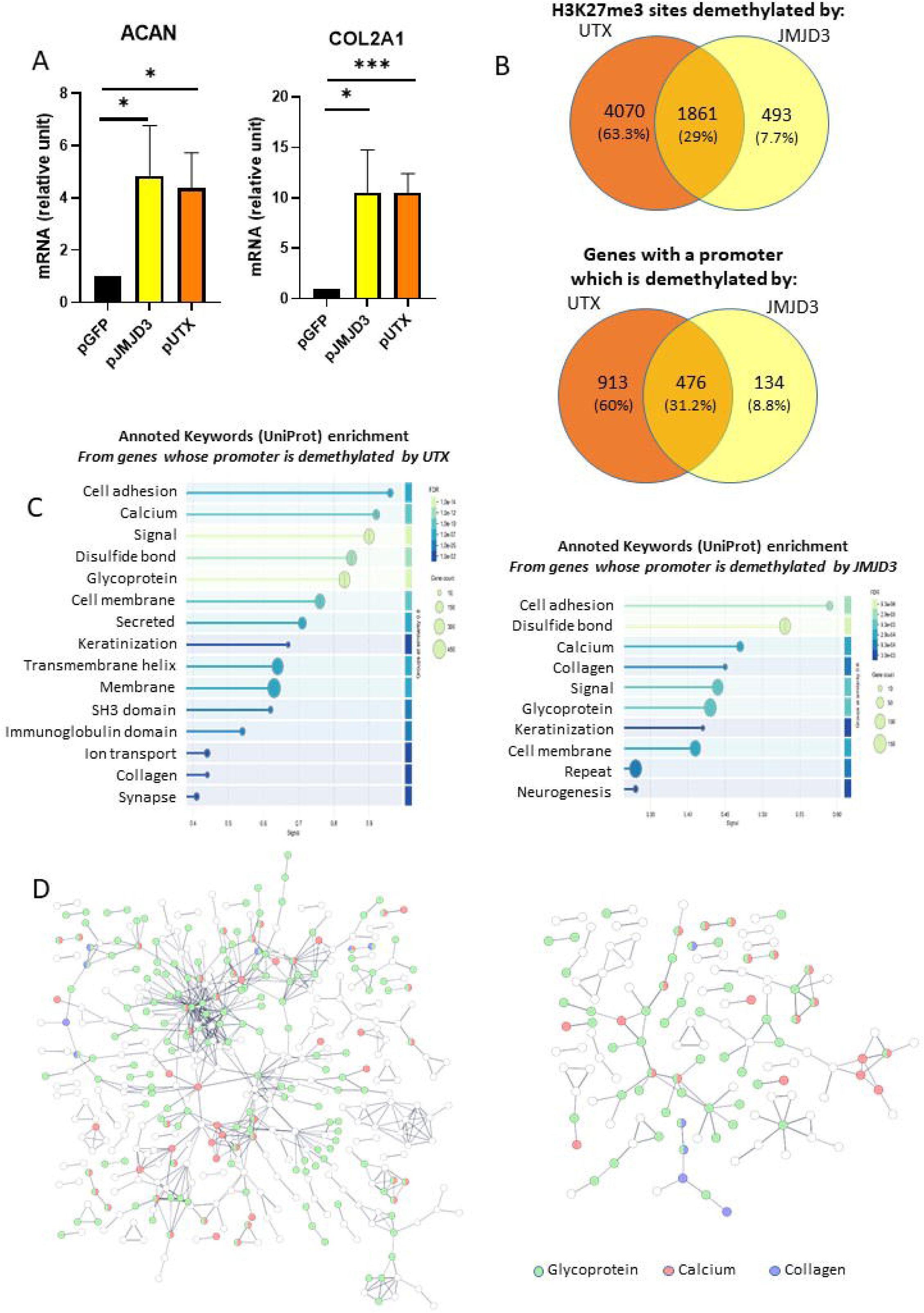
JMJD3 and UTX enhance chondrogenesis and share common target genes. hBM-MSCs were nucleofected with expression vectors for GFP (pCtrl), JMJD3 (pJMJD3), or UTX (pUTX). After nucleofection, cells were cultured in alginate beads in chondrogenic medium for 7 days. (A) mRNA was extracted, and the relative expression levels of ACAN and COL2A1 were determined by RT-PCR. Data are presented as mean ± SEM (*n* = 5). *: pvalue < 0.05; ***: pvalue < 0.001. (B) ChIP-Seq was performed to identify promoters with H3K27me3 peaks present in pGFP-transfected cells but absent in pJMJD3- or pUTX-transfected cells. Venn diagrams comparing regions demethylated by pUTX or pJMJD3 relative to pGFP are shown. (C, D) Enrichment analysis of annotated keywords (UNIPROT) and protein-protein interaction (PPI) networks were generated using the STRING database.

### JMJD3 and UTX partially share overlapping targets

To gain deeper insights into the functions of JMJD3 and UTX during chondrogenesis and to investigate whether they might act redundantly by targeting the same genomic loci, we analyzed ChIP-Seq data from three distinct conditions: “GFP” (hBM-MSCs transiently transfected with a GFP expression vector as a control before embedding in alginate beads and cultured in chondrogenic medium for 7 days), “JMJD3” (cells transfected with a JMJD3 expression vector before inducing chondrogenesis), and “UTX” (cells transfected with a UTX expression vector before inducing chondrogenesis). We focused on genes whose promoters were H3K27me3 undermethylated upon JMJD3 or UTX overexpression, i.e. genes with H3K27me3 enrichment near their promoters in the control condition (“GFP”) but not in either the “JMJD3” or “UTX” conditions. ChIP-Seq revealed 2 356 H3K27me3 peaks at the promoters of 610 genes in the “GFP” condition that were absent in the “JMJD3” condition but present in “GFP” condition. Similarly, 5 937 H3K27me3 peaks corresponding to 1 389 genes were present in the “GFP” condition but absent in the “UTX” condition. Comparative analysis of these datasets (Figure 5B) identified 1 861 H3K27me3 sites (29%) that were erased following both JMJD3 and UTX overexpression, corresponding to 476 genes (31%). This overlap indicates that JMJD3 and UTX share numerous common targets. Notably, the genes with H3K27me3 sites demethylated at their promoters upon overexpression of either demethylase were enriched for genes encoding proteins annotated by UniProt as “collagen,” “glycoprotein,” or “calcium”, suggesting potential redundant functions and overlapping targets for JMJD3 and UTX (Figures 5C and 5D).

### JMJD3 and UTX enhanced in vivo cartilage formation

To evaluate the potential application of demethylase overexpression for tissue engineering, given its enhancement of chondrogenesis in vitro, we conducted in vivo experiments. Alginate beads containing hBM-MSCs transiently transfected with expression vectors for JMJD3, UTX, or GFP (as a control) and subjected to 7 days of in vitro chondrogenic differentiation were subcutaneously implanted into nude mice for four weeks (Figure 6A).

**Figure 6:**
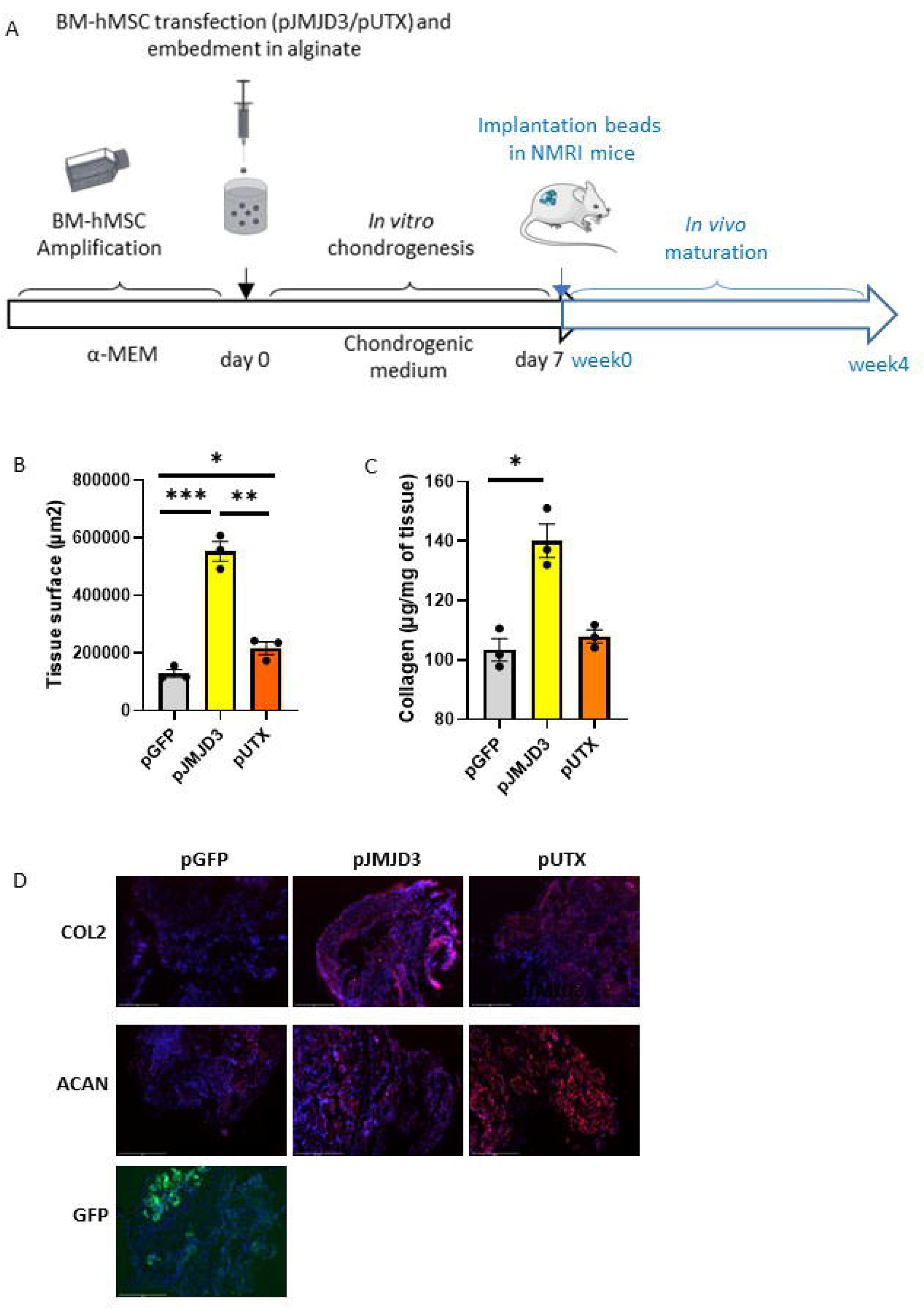
JMJD3 and UTX enhance in vivo cartilage formation. After transfection with expression vectors for GFP (pGFP), JMJD3 (pJMJD3), or UTX (pUTX), hBM-MSCs were embedded in alginate beads and cultured in chondrogenic medium for 7 days. The beads were then implanted subcutaneously into mice for 4 weeks. (A) Design of the experiment. (B) Cartilage disk surface area was measured. (C) Collagen content was quantified using the Sircol assay. Data are expressed as mean ± SEM (*n* = 3). * = pvalue < 0.05 ** = pvalue < 0.01, *** = pvalue < 0.001. (D) Immunohistofluorescence staining was performed to detect type II collagen and aggrecan expression in the cartilage tissue. Fluorescence emitted by GFP is also shown.

Cartilage discs containing cells having overexpressed JMJD3 were significantly larger than those containing GFP-transfected control cells (Figure 6B). Similarly, discs from UTX-overexpressing cells were larger than the control group, though smaller than those from JMJD3-overexpressing cells. We then quantified the total collagen content within the discs. As shown in figure 6C, JMJD3 overexpression significantly increased collagen content compared to the GFP control. Interestingly, this increase in collagen was attributed, at least in part, to elevated type II collagen expression (Figure 6D). UTX overexpression also enhanced type II collagen expression, albeit to a lesser extent than JMJD3. Additionally, ACAN expression appeared to be elevated following both JMJD3 and UTX overexpression. These results confirm that JMJD3 and UTX promote chondrogenesis, with JMJD3 exhibiting a more pronounced effect on cartilage formation and extracellular matrix production *in vivo*.

## Discussion

Genetically modified MSCs represent a promising strategy for cartilage regeneration. Various transgenes have been tested to stimulate articular cartilage restoration, yet none have fully optimized chondrogenesis. A deeper understanding of the molecular mechanisms involved, particularly epigenetic regulation, may enhance cartilage repair outcomes. In this study, we investigated the regulation and role of the histone demethylases JMJD3 and UTX during *in vitro* chondrogenesis of human bone marrow-derived mesenchymal stem cells. Our findings demonstrate that while only JMJD3 (but not UTX) is essential for chondrogenesis, both JMJD3 and UTX can demethylate similar genomic loci in genome, favoring the expression of chondrogenic signature genes and improving *in vitro* chondrogenesis. Consequently, these H3K27me3 demethylases emerge as potential candidates for improving gene-modified MSC therapy in cartilage tissue engineering.

We utilized primary hBM-MSCs from osteoarthritis donors, differentiated towards a chondrocyte phenotype over seven days in a 3D culture system using alginate beads and TGF-β3. This model, widely recognized for inducing in vitro chondrogenesis^4,15^, allowed us to profile H3K27me3 and H3K4me3 using ChIP-Seq. At the time of the experiments (7 days), we did not observe a reduction in H3K27me3 enrichment at chondrogenic gene loci. However, over 10% of H3K27me3 sites were demethylated during differentiation, underscoring the pivotal role of H3K27me3 demethylation in this process. These results align with Herlofsen et al. (2013), who, using a different differentiation protocol (TGF-β1 and BMP2), similarly reported persistent H3K27me3 at the COL2A1 locus but noted increased H3K4me3 near its transcription start site (TSS) ^16^.

To further elucidate the role of H3K27me3 demethylation, we induced chondrogenesis in the presence of GSK-J4, the only known inhibitor of KDM6 family demethylases (JMJD3 and UTX), and analyzed gene expression and H3K27me3 profiles. GSK-J4 treatment reduced ACAN and COL2A1 expression, correlating with additional H3K27me3 peaks at their promoters, suggesting that GSK-J4 impedes chondrogenesis by inhibiting H3K27me3 demethylase activity. Similarly, Yapp et al. showed that GSK-J4 does not affect adipogenic or osteogenic gene expression, but reduces the expression of *SOX9*, *COL2A1*, *ACAN* and *COL10A1* and it increased *COL1A1* expression during *in vitro* chondrogenesis of human bone marrow mesenchymal stem cells cultured in monolayer^17^. They also observed that GSK-J4 treatment decreases the glycosaminoglycan content in cartilage discs, and disorganizes collagen fibers.

Similarly, Yapp et al. demonstrated that GSK-J4 does not affect adipogenic or osteogenic gene expression but significantly reduces the expression of SOX9, COL2A1, ACAN, and COL10A1 during in vitro chondrogenesis of human bone marrow-derived mesenchymal stem cells cultured in monolayer conditions. Additionally, they observed that GSK-J4 treatment decreases the glycosaminoglycan content in cartilage discs derived from MSCs cultured in polycarbonate Transwell filters with chondrogenic medium in the lower chamber, along with the presence of disorganized collagen fibers.

We identified several genes, including ACAN, HEY2, and LEF1, as direct targets of JMJD3 or UTX. These genes, key players in chondrogenesis^18,19^, were downregulated by GSK-J4 and exhibited new H3K27me3 peaks at their promoters after GSK-J4 treatment. Gene ontology analysis revealed that GSK-J4-downregulated genes are involved in “chondrocyte differentiation,” “skeletal system development,” “TGF-β signaling,” and “Wnt signaling,” aligning with the established roles of H3K27me3 demethylases. Notably, JMJD3 deletion in mice delays endochondral ossification^5^. Additionally, Wnt signaling, which is a critical chondrogenic regulator, mediated the function of KDM6A/KDM6B during early human embryonic stem cell differentiation^20^.

GSK-J4 is described as an inhibitor of the KDM6 family of H3K27me3 demethylases (*i.e*. JMJD3 and UTX) able to impaired the reduction of H3K27me3 observed during chondrogenesis ^14,17^, but it has also been shown that GSK-J4 is able to target the KDM5 family of H3K4me3 demethylases ^21^. So, to distinguish the specific contributions of JMJD3 and/or UTX, we performed siRNA-mediated knockdowns. In agreement with previous published study ^17^, the knock-down of JMJD3 inhibited hBM-MSC chondrogenic differentiation by reducing *ACAN*, *COL2A1*, *SOX5* and *SOX6* expression. At contrary, UTX knockdown had no significant effect, suggesting that the impact of GSK-J4 comes mainly from the inhibition of JMJD3. This may be due to the upregulation of JMJD3 during chondrogenesis, unlike UTX, which remains at basal levels.

Interestingly, overexpression of both JMJD3 and UTX enhanced ACAN and COL2A1 expression, with H3K27me3 profiling indicating overlapping target loci. The fact that UTX knock-down had no impact on chondrogenesis in contrast to its forced expression may be explained by its transcriptional regulation. Indeed, JMJD3 is generally considered to be induced by stimuli while UTX is expressed at basal level whatever the conditions^12,22,23^. Herein, in agreement with a previous study^17^, only JMJD3 was upregulated during *in vitro* chondrogenesis. Consequently, the knock-down of UTX may have no effect on chondrogenesis because UTX stays at a basal expression level. Therefore, although UTX and JMJD3 are partially redundant, it appears that they differ in spatio-temporal expression leading to different activity during mesenchymal differentiation. A similar regulatory mechanism has been proposed between EZH2 and EZH1, the both enzymes able to catalyze H3K27 trimethylation ^24^.

Supporting Yapp’s hypothesis^17^, we demonstrated that transient overexpression of JMJD3 and UTX improves cartilage formation. Cartilage discs derived from hBM-MSCs transfected with JMJD3 or UTX were larger and richer in collagen than control discs, demonstrating the potential of enhancing H3K27me3 demethylase activity to favor chondrogenesis. However, the expression of these histone demethylases will require to be manipulated with precaution, as they can exacerbate osteoarthritis in mature chondrocytes. ^6,8,25^. This suggests that the forced expression of JMJD3 or UTX will have to be transient. The expression of these demethylases must be tightly regulated. Therefore, transient upregulation during early chondrogenic differentiation may be optimal.

In conclusion, our study highlights the critical role of JMJD3 and UTX in chondrogenesis. While JMJD3 is indispensable, both demethylases enhance chondrogenesis and cartilage formation when overexpressed. These findings provide valuable insights into the epigenetic control of chondrogenesis and suggest JMJD3 and UTX as promising targets for gene-modified MSC therapy in cartilage tissue engineering.

## Acknowledgements

We thank Sylvain Leclercq (Service orthopédique de la Clinique Saint-Martin, Caen France) for the human tissue samples. We also thank OncoModels platform (UNICAEN, Caen) for assistance for in vivo experiments, and Proteogen platform for providing equipments required for transcriptomics analysis.

## Fundings

This study was supported by the Agence Nationale de la Recherche [ANR-15-CE14-0002-01], Région Normandie, European Union [FEDER/FSE 2014-2020 – 16E00779/16P03685], and Société Française de Rhumatologie (SFR). Lyess Allas was recipient of a doctoral allocation from University of Caen Normandie.

## Author contributions: CRediT

LA and CB designed the study. LA acquired and analysed the majority of the data. LA, JAL, QR, AJ, EL, ML, SB participated to in vitro experiments. LA and AV performed in vivo experiments. JAL performed –omics analysis and visualization. LA and CB did statistical analysis. LA, JAL, KB and CB interpreted the data. CB and KB administered the project. CB supervised the project. CB obtained fundings. LA and CB drafted the original manuscript. JAL brought substantial revision of the primary draft. All authors provided revisions and approved the final version of the manuscript.

## Conflicts of interest

Authors have nothing to declare

## Declaration of generative AI and AI-assisted technologies in the writing process

During the preparation of this work the authors used ChatGPT in order to improve the readability and language of the manuscript. After using this tool, the authors reviewed and edited the content as needed and take full responsibility for the content of the published article.

## Notes

### Competing Interest Statement

The authors have declared no competing interest.

## References

1. Brittberg, M. et al. Treatment of deep cartilage defects in the knee with autologous chondrocyte transplantation. N. Engl. J. Med. 331, 889–895 (1994).

2. Duval, E. et al. Hypoxia-inducible factor 1alpha inhibits the fibroblast-like markers type I and type III collagen during hypoxia-induced chondrocyte redifferentiation: hypoxia not only induces type II collagen and aggrecan, but it also inhibits type I and type III collagen in the hypoxia-inducible factor 1alpha-dependent redifferentiation of chondrocytes. Arthritis Rheum. 60, 3038–3048 (2009).

3. Im, G.-I. & Shin, K.-J. Epigenetic approaches to regeneration of bone and cartilage from stem cells. Expert Opin. Biol. Ther. 15, 181–193 (2015).

4. Baugé, C. & Boumédiene, K. Use of Adult Stem Cells for Cartilage Tissue Engineering: Current Status and Future Developments. Stem Cells Int. 2015, e438026 (2015).

5. Zhang, F. et al. JMJD3 promotes chondrocyte proliferation and hypertrophy during endochondral bone formation in mice. J. Mol. Cell Biol. 7, 23–34 (2015).

6. Lian, W.-S. et al. Histone H3K27 demethylase UTX compromises articular chondrocyte anabolism and aggravates osteoarthritic degeneration. Cell Death Dis. 13, 538 (2022).

7. Jun, Z. et al. Jumonji domain containing-3 (JMJD3) inhibition attenuates IL-1β-induced chondrocytes damage in vitro and protects osteoarthritis cartilage in vivo. Inflamm. Res. Off. J. Eur. Histamine Res. Soc. Al 69, 657–666 (2020).

8. Jin, Y. et al. Histone demethylase JMJD3 downregulation protects against aberrant force-induced osteoarthritis through epigenetic control of NR4A1. Int. J. Oral Sci. 14, 1–14 (2022).

9. Duval, E. et al. Molecular mechanism of hypoxia-induced chondrogenesis and its application in in vivo cartilage tissue engineering. Biomaterials 33, 6042–6051 (2012).

10. Szklarczyk, D. et al. The STRING database in 2023: protein-protein association networks and functional enrichment analyses for any sequenced genome of interest. Nucleic Acids Res. 51, D638–D646 (2023).

11. Aury-Landas, J. et al. Anti-inflammatory and chondroprotective effects of the S-adenosylhomocysteine hydrolase inhibitor 3-Deazaneplanocin A, in human articular chondrocytes. Sci. Rep. 7, (2017).

12. Agger, K. et al. UTX and JMJD3 are histone H3K27 demethylases involved in HOX gene regulation and development. Nature 449, 731–734 (2007).

13. Allas, L., Rochoux, Q., Leclercq, S., Boumédiene, K. & Baugé, C. Development of a simple osteoarthritis model useful to predict in vitro the anti-hypertrophic action of drugs. Lab. Investig. J. Tech. Methods Pathol. (2019) doi:10.1038/s41374-019-0303-0.

14. Kruidenier, L. et al. A selective jumonji H3K27 demethylase inhibitor modulates the proinflammatory macrophage response. Nature 488, 404–408 (2012).

15. Hammad, M. et al. Hypoxia Differentially Affects Chondrogenic Differentiation of Progenitor Cells from Different Origins. Int. J. Stem Cells 16, 304–314 (2023).

16. Herlofsen, S. R. et al. Genome-wide map of quantified epigenetic changes during in vitro chondrogenic differentiation of primary human mesenchymal stem cells. BMC Genomics 14, 105 (2013).

17. Yapp, C., Carr, A. J., Price, A., Oppermann, U. & Snelling, S. J. B. H3K27me3 demethylases regulate in vitro chondrogenesis and chondrocyte activity in osteoarthritis. Arthritis Res. Ther. 18, 158 (2016).

18. Canalis, E., Yu, J., Singh, V., Mocarska, M. & Schilling, L. NOTCH 2 Sensitizes the Chondrocyte to the Inflammatory Response of Tumor Necrosis Factor α. J. Biol. Chem. 0, (2023).

19. Schizas, N. P. et al. Inhibition versus activation of canonical Wnt-signaling, to promote chondrogenic differentiation of Mesenchymal Stem Cells. A review. Orthop. Rev. 13, (2021).

20. Jiang, W., Wang, J. & Zhang, Y. Histone H3K27me3 demethylases KDM6A and KDM6B modulate definitive endoderm differentiation from human ESCs by regulating WNT signaling pathway. Cell Res. 23, 122–130 (2013).

21. Heinemann, B. et al. Inhibition of demethylases by GSK-J1/J4. Nature 514, E1–2 (2014).

22. Van der Meulen, J., Speleman, F. & Van Vlierberghe, P. The H3K27me3 demethylase UTX in normal development and disease. Epigenetics 9, 658–668 (2014).

23. Arcipowski, K. M., Martinez, C. A. & Ntziachristos, P. Histone demethylases in physiology and cancer: a tale of two enzymes, JMJD3 and UTX. Curr. Opin. Genet. Dev. 36, 59–67 (2016).

24. Camilleri, E. T. et al. Loss of histone methyltransferase Ezh2 stimulates an osteogenic transcriptional program in chondrocytes but does not affect cartilage development. J. Biol. Chem. 293, 19001–19011 (2018).

25. Lian, W.-S. et al. Inhibition of histone lysine demethylase 6A promotes chondrocytic activity and attenuates osteoarthritis development through repressing H3K27me3 enhancement of Wnt10a. Int. J. Biochem. Cell Biol. 158, 106394 (2023).

