## Supplementary material for "Epigenetic Regulation of Chondrogenesis: JMJD3 and UTX as Key Targets for Gene-Modified Mesenchymal Stem Cell Therapy in Cartilage Tissue Engineering": Suppl table 1

**Suppl table 1 : List of DEG**

| ProbeName | FC ([MSC] vs Zero) | Regulation ([I | GeneSymbol |
|---|---|---|---|
| A_33_P3421163 | 1.652419 | up | TP53INP1 |
| (+)E1A_r60_a22 | 8.974959 | up | |
| A_23_P118392 | 1.6158189 | up | RASD1 |
| A_23_P118246 | -1.7651154 | down | GINS2 |
| A_24_P943894 | -2.0480661 | down | SCUBE3 |
| A_23_P132738 | -3.2171526 | down | CRYGS |
| A_23_P10121 | -1.5934201 | down | SFRP1 |
| A_23_P117852 | -2.4492047 | down | KIAA0101 |
| A_33_P3268838 | 1.9713067 | up | CPEB1 |
| A_21_P0000504 | 1.9188547 | up | RNU6ATAC |
| A_23_P427703 | 3.7832692 | up | MT1L |
| (+)E1A_r60_a107 | -2.2715554 | down | |
| (+)E1A_r60_a135 | -2.188322 | down | |
| (+)E1A_r60_3 | 3.2951138 | up | |
| A_33_P3339212 | -2.7086153 | down | TRIP13 |
| A_33_P3332135 | -2.2408178 | down | PHOSPHO1 |
| A_33_P3277407 | 1.7901449 | up | KMT2E |
| A_23_P258493 | -2.501572 | down | LMNB1 |
| A_22_P00009778 | 2.3232102 | up | |
| A_33_P6817068 | -1.5548127 | down | NDUFA6-AS1 |
| A_23_P25706 | 1.7889777 | up | CLMN |
| (+)E1A_r60_a97 | 2.0847874 | up | |
| A_24_P297539 | -3.7035918 | down | UBE2C |
| A_24_P192301 | -2.716889 | down | SEMA3A |
| A_24_P226069 | -2.0613692 | down | FGFBP2 |
| A_23_P122007 | 1.5079508 | up | C5orf30 |
| A_23_P13907 | -2.4059074 | down | IGF1 |
| (+)E1A_r60_n9 | -6.5770783 | down | |
| A_33_P3426943 | -2.9293294 | down | LOC151484 |
| A_32_P74409 | 1.6400545 | up | C11orf96 |
| A_33_P3814721 | -2.5447404 | down | INSC |
| A_33_P3235147 | -1.9993324 | down | DLX5 |
| A_33_P3306327 | 1.5230261 | up | |
| A_23_P14986 | -3.7361865 | down | HSD11B2 |
| (+)E1A_r60_a104 | 3.261466 | up | |
| A_23_P256948 | 2.1347187 | up | MSC |
| (+)E1A_r60_n11 | -1.9353248 | down | |
| A_23_P324327 | -2.6806011 | down | GPRC5B |
| A_23_P41804 | -2.2441905 | down | NKD2 |
| A_33_P3387601 | -1.7144762 | down | KLKP1 |
| A_23_P33326 | 2.1756983 | up | ADRA1B |
| A_33_P3322804 | -2.108775 | down | NTRK2 |
| A_23_P118815 | -3.042028 | down | BIRC5 |
| A_23_P88522 | 2.6421754 | up | NMB |
| A_23_P106617 | -2.1385374 | down | WFDC1 |
| A_23_P131935 | -1.6946663 | down | FERMT1 |
| A_21_P0000418 | 1.7380116 | up | SNORD114-21 |
| A_24_P34155 | -1.6870791 | down | RUNX1 |
| A_33_P3222380 | 1.612025 | up | AHNAK2 |

| | | | |
|---|---|---|---|
| A_23_P41424 | -1.523455 | down | SLC39A8 |
| A_23_P103720 | -1.5846052 | down | AGMAT |
| A_23_P7313 | -1.9897714 | down | SPP1 |
| A_33_P3231923 | 1.5589771 | up | LOC101927285 |
| A_23_P149626 | -2.045746 | down | PLEKHG5 |
| A_33_P3341490 | -1.7125988 | down | KRT42P |
| A_23_P71328 | -1.7946984 | down | MATN2 |
| A_33_P3238997 | 1.844409 | up | AGFG1 |
| A_33_P3409086 | -2.4749556 | down | S100A1 |
| A_23_P121064 | 2.7474241 | up | PTX3 |
| A_33_P3236868 | 3.3513103 | up | MT1X |
| A_33_P3295343 | -7.917364 | down | CLEC3A |
| A_23_P259594 | -1.8512341 | down | AKAP7 |
| A_33_P3239512 | 1.8681624 | up | ZNF836 |
| A_19_P00316344 | 1.5901246 | up | lnc-C9orf69-2 |
| A_33_P3308332 | -2.021994 | down | PLEKHB1 |
| A_33_P3375934 | 2.172199 | up | NAMPT |
| A_33_P3227793 | -1.5338856 | down | CGREF1 |
| A_24_P68908 | 3.8249757 | up | LOC344887 |
| A_24_P33156 | -1.6835511 | down | AFMID |
| A_23_P62081 | -1.5446848 | down | SCG5 |
| A_23_P32414 | -1.6360377 | down | STK26 |
| A_23_P139820 | 1.7751114 | up | SLC11A2 |
| A_23_P383009 | -2.6147912 | down | IGFBP5 |
| A_21_P0000406 | 1.6441603 | up | SNORD114-9 |
| A_23_P369994 | 1.9547948 | up | DCLK1 |
| A_23_P111995 | -1.6801388 | down | LOXL2 |
| A_32_P167239 | 2.3066328 | up | AFAP1L1 |
| A_23_P53176 | -1.5511059 | down | FOLR1 |
| A_23_P32253 | 1.564841 | up | NFIL3 |
| A_33_P3379106 | 2.3241956 | up | LINC00922 |
| A_23_P254688 | -2.7297964 | down | TMEM108 |
| A_24_P941167 | 1.9387273 | up | APOL6 |
| A_33_P3398156 | -1.6312131 | down | CYS1 |
| A_23_P213893 | -1.9910079 | down | MEGF10 |
| A_24_P212539 | 1.6585654 | up | GALM |
| A_32_P342064 | 2.1558712 | up | FTH1 |
| A_19_P00318759 | -1.5367509 | down | APCDD1L-AS1 |
| A_24_P10137 | -2.682681 | down | RGCC |
| A_23_P342138 | 1.841086 | up | ADAMTSL1 |
| A_33_P3335371 | -2.1343076 | down | MAML3 |
| A_23_P30254 | -1.9440174 | down | PLK2 |
| A_23_P55448 | -2.1490767 | down | KRT12 |
| A_24_P130363 | -1.9291517 | down | LDLRAD4 |
| A_24_P273253 | 1.6379284 | up | AHNAK2 |
| A_23_P138507 | -3.0815277 | down | CDK1 |
| A_33_P3339531 | -2.073449 | down | CHADL |
| A_23_P35617 | -1.5175923 | down | PLCE1 |
| A_23_P82990 | -2.4068239 | down | OGN |
| A_24_P224727 | 2.2424707 | up | CEBPA |
| A_33_P3410599 | -1.5706533 | down | FAM46A |

| | | | |
|---|---|---|---|
| A_33_P3376463 | 1.7240683 | up | NR2C2 |
| A_24_P133253 | 1.9270607 | up | KITLG |
| A_23_P206684 | -1.8348253 | down | WWP2 |
| A_24_P185854 | 1.7219236 | up | DMD |
| A_23_P15798 | -2.7225246 | down | KRTAP4-12 |
| A_21_P0005976 | 1.5245316 | up | lnc-ASAP1-1 |
| A_23_P318904 | -2.0147336 | down | SERTAD4 |
| A_23_P317729 | -2.8327262 | down | SP7 |
| A_23_P69908 | 2.0067108 | up | GLRX |
| A_33_P3343290 | 2.0784774 | up | PPTC7 |
| A_23_P34915 | 4.2299514 | up | ATF3 |
| A_32_P203404 | -2.972479 | down | FAM69C |
| A_33_P3338698 | -3.2107322 | down | IHH |
| A_23_P74278 | -2.0003104 | down | PDE4B |
| A_33_P3273298 | 1.8628681 | up | ZNF813 |
| A_24_P253003 | -2.554021 | down | WNT11 |
| A_23_P398294 | 1.7197443 | up | HIP1R |
| A_33_P3391076 | 1.6947967 | up | MBNL2 |
| A_23_P407840 | -2.131316 | down | FNDC1 |
| A_24_P286951 | -4.3840675 | down | PLEKHS1 |
| A_33_P3331746 | -5.698503 | down | ST6GAL2 |
| A_23_P44466 | 2.0877714 | up | CCDC102B |
| A_22_P00002306 | -2.703651 | down | MIR210HG |
| A_33_P3210085 | -1.8383814 | down | NET1 |
| A_24_P381494 | 1.7693995 | up | SLC11A2 |
| A_33_P3211929 | -2.3166168 | down | RCOR2 |
| A_33_P3400763 | 1.9955107 | up | PLIN4 |
| A_32_P37867 | 1.7728349 | up | KIAA1644 |
| A_22_P00013487 | -1.8855842 | down | lnc-RP11-334E6.3.1-2 |
| A_21_P0011028 | 1.6443799 | up | LOC102724910 |
| A_24_P183664 | -1.6952221 | down | TRIL |
| A_24_P216654 | 1.7435976 | up | SOAT1 |
| A_33_P3382924 | -1.5336068 | down | SPARC |
| A_33_P3317460 | -1.6722713 | down | |
| A_33_P3334630 | -3.129482 | down | PLP1 |
| A_23_P59613 | -1.707076 | down | FZD9 |
| A_23_P167030 | -3.0295172 | down | PTH1R |
| A_23_P43164 | -1.7825164 | down | SULF1 |
| A_33_P3376249 | -1.7453945 | down | S100A2 |
| A_23_P106002 | 1.585836 | up | NFKBIA |
| A_23_P33196 | -1.6096483 | down | COL5A2 |
| A_21_P0000507 | -2.5555408 | down | SNAR-B2 |
| A_24_P321525 | -2.8870404 | down | RERG |
| A_23_P74609 | 2.0508974 | up | G0S2 |
| A_23_P69738 | -1.699325 | down | RASL11B |
| A_33_P3311775 | -1.766298 | down | ZNF789 |
| A_22_P00005618 | -2.2260392 | down | |
| A_33_P3312509 | -2.2238393 | down | ADAMTSL2 |
| A_33_P3246838 | -2.0666058 | down | |
| A_23_P62335 | -1.7460243 | down | TMLHE |
| A_22_P00007123 | -2.388703 | down | |

| | | | |
|---|---|---|---|
| A_33_P3229156 | -1.5610328 | down | SLC17A9 |
| A_21_P0014771 | 1.9426671 | up | |
| A_33_P3807062 | -2.066917 | down | HJURP |
| A_24_P314179 | -1.709673 | down | ETS2 |
| A_23_P153676 | 1.8452277 | up | TLE2 |
| A_23_P56356 | -1.9076135 | down | PLB1 |
| A_33_P3413958 | -1.7817045 | down | OR7E47P |
| A_23_P432947 | -2.7464008 | down | GREM1 |
| A_19_P00315581 | 1.5389221 | up | LINC01122 |
| A_32_P51237 | -1.9089172 | down | KANK4 |
| A_23_P431971 | -2.0878527 | down | CRTAC1 |
| A_33_P3272231 | 1.6779113 | up | MFSD2A |
| A_24_P179044 | 2.151919 | up | SNX9 |
| A_23_P383227 | -2.259042 | down | S100A1 |
| A_33_P3267186 | 2.098451 | up | ATP2B1 |
| A_32_P208403 | -1.9881378 | down | GNG2 |
| A_33_P3369098 | -2.3698812 | down | MYL10 |
| A_33_P3315314 | 3.4239643 | up | MT1HL1 |
| A_33_P3327587 | -1.7985157 | down | LOC100127904 |
| A_23_P40295 | -2.044884 | down | LAMP5 |
| A_21_P0001085 | -1.7560272 | down | LOC100130417 |
| A_19_P00319409 | -2.7767866 | down | ERVMER34-1 |
| A_24_P406060 | -1.8853822 | down | RNF144B |
| A_21_P0008540 | -1.8725497 | down | lnc-MAP3K9-3 |
| A_23_P126844 | -1.6366007 | down | TNFRSF25 |
| A_23_P215956 | -1.5829742 | down | MYC |
| DCP_22_0 | 9.048078 | up | |
| A_24_P318656 | 2.0660405 | up | ITGB3 |
| A_19_P00323082 | -1.9613239 | down | H19 |
| A_23_P79622 | -1.6171749 | down | FKBP7 |
| A_24_P68247 | 1.5879569 | up | TRIM4 |
| A_33_P3268472 | -2.3200972 | down | CTSC |
| A_23_P205713 | -1.6586232 | down | STXBP6 |
| A_23_P102058 | -2.330327 | down | MATN3 |
| A_23_P120243 | -3.0940893 | down | HOXD1 |
| A_32_P196263 | -2.1501913 | down | ADAMTS9 |
| A_23_P363778 | -1.8411065 | down | FRZB |
| A_24_P157926 | 1.8127707 | up | TNFAIP3 |
| DCP_22_6 | 9.443785 | up | |
| A_33_P3376095 | 1.9878775 | up | SYPL2 |
| A_23_P300600 | -1.6864964 | down | NEFH |
| A_23_P68031 | -2.1865587 | down | STAT4 |
| A_33_P3347452 | 1.981046 | up | RPS6KA2 |
| A_23_P48835 | -2.0624123 | down | KIF23 |
| A_24_P753454 | -1.6372671 | down | NDUFA6-AS1 |
| A_23_P70007 | -2.2827046 | down | HMMR |
| A_23_P206724 | 4.341748 | up | MT1E |
| A_23_P500501 | -3.3670604 | down | FGFR3 |
| A_23_P137573 | -2.0551069 | down | LEFTY2 |
| A_33_P3268313 | -2.5362787 | down | PGAM2 |
| A_33_P3298024 | 1.649117 | up | ABCC3 |

| | | | |
|---|---|---|---|
| DCP_22_4 | 9.134026 | up | |
| A_21_P0000508 | -2.6012263 | down | SNAR-D |
| A_21_P0011496 | 2.4448562 | up | CES1 |
| A_23_P68610 | -1.8460907 | down | TPX2 |
| A_23_P144549 | -4.276504 | down | IBSP |
| A_33_P3391375 | 1.6070416 | up | LANCL3 |
| A_19_P00320067 | -1.720521 | down | LOC100130417 |
| A_23_P87072 | -3.5639663 | down | PANX3 |
| A_23_P47148 | -1.6503834 | down | NOX4 |
| A_21_P0005787 | 2.1868968 | up | LOC101927815 |
| A_23_P40108 | -5.434459 | down | COL9A3 |
| A_23_P104563 | 1.696495 | up | CPT1A |
| A_23_P114883 | -1.9058226 | down | FMOD |
| A_23_P207507 | 1.9742265 | up | ABCC3 |
| A_33_P3253249 | 2.1913505 | up | FTH1 |
| A_24_P140475 | 1.6488314 | up | SORBS2 |
| A_33_P3449097 | 1.5064813 | up | TSPAN10 |
| A_23_P48669 | -1.7143297 | down | CDKN3 |
| A_24_P258846 | -1.6592423 | down | NFATC1 |
| A_23_P41629 | -2.6286228 | down | ADAMTS16 |
| A_33_P3590259 | -7.049741 | down | CXCL14 |
| A_33_P3247057 | -3.6005754 | down | SMIM5 |
| A_33_P3279590 | -3.2860255 | down | OGN |
| A_33_P3245178 | 4.035224 | up | BEX2 |
| A_24_P286114 | 2.2175918 | up | SLC1A3 |
| A_22_P00021205 | -2.5448132 | down | HAGLROS |
| A_22_P00010235 | 3.4634519 | up | MT1B |
| A_23_P138435 | 1.7010924 | up | ZMIZ1 |
| A_33_P3265270 | 1.9634409 | up | SLC17A5 |
| A_33_P3424132 | -1.7611881 | down | LINC00961 |
| A_21_P0014577 | -2.2770963 | down | LOC100506869 |
| A_23_P371266 | -1.8103201 | down | DNM3 |
| A_33_P3246418 | -1.882626 | down | MDFI |
| A_24_P139901 | 1.7039294 | up | GYPC |
| A_32_P25357 | -1.969251 | down | CDH15 |
| A_23_P111311 | 1.5880468 | up | AKAP12 |
| A_23_P99063 | -1.7210608 | down | LUM |
| A_21_P0001177 | -1.5671433 | down | lnc-FAM72C-2 |
| A_33_P3335590 | -2.3918204 | down | CCDC64B |
| A_23_P52017 | -2.5209954 | down | ASPM |
| A_21_P0000419 | 1.787493 | up | SNORD114-22 |
| A_23_P4714 | -3.377358 | down | MIA |
| A_23_P65757 | -1.8416226 | down | CCNB2 |
| A_33_P3283480 | -2.184844 | down | CTSC |
| A_21_P0009476 | 2.82912 | up | lnc-KATNAL2-4 |
| A_33_P3257678 | -4.6536393 | down | HIST2H3A |
| A_33_P3670415 | 1.6030469 | up | NAT8L |
| A_32_P108655 | -1.6497557 | down | AK4 |
| A_23_P41114 | 2.243246 | up | CSTA |
| A_22_P00014719 | -2.115151 | down | LOC101929726 |
| A_33_P3350726 | 1.6275526 | up | PPARG |

| | | | |
|---|---|---|---|
| A_24_P126210 | 2.7870111 | up | MRAP |
| A_23_P48676 | -1.5623227 | down | PYGL |
| A_23_P375 | -1.9881757 | down | CDCA8 |
| A_23_P70307 | -4.8499355 | down | SMOC2 |
| A_23_P42322 | -2.8179874 | down | COL11A2 |
| A_33_P3333282 | -2.7342763 | down | FGF11 |
| A_33_P3364864 | 2.6576314 | up | NAMPT |
| DCP_22_7 | 8.539384 | up | |
| A_32_P105549 | -2.310197 | down | ANXA8L1 |
| A_33_P3234764 | -2.3307307 | down | |
| A_24_P321766 | -2.1722677 | down | SERPINA5 |
| A_33_P3393766 | -1.7562927 | down | C17orf96 |
| A_23_P63209 | 2.6981313 | up | HSD11B1 |
| A_23_P26847 | -1.7236811 | down | SOX9 |
| A_33_P3289113 | 1.7336009 | up | COX11 |
| A_24_P152968 | 2.3042178 | up | AKR1C1 |
| A_23_P65518 | -1.5188963 | down | DACT1 |
| A_22_P00013705 | 1.8412241 | up | ZFPM2-AS1 |
| DCP_22_2 | 9.116553 | up | |
| A_23_P252306 | -2.751872 | down | ID1 |
| A_32_P59010 | 2.287197 | up | TNFAIP8L3 |
| A_33_P3345534 | -2.3364587 | down | KRT14 |
| A_21_P0000511 | -2.6034384 | down | SNAR-H |
| A_23_P158096 | -1.5492457 | down | COL27A1 |
| A_33_P3235262 | -1.6821489 | down | PIP5KL1 |
| A_23_P60146 | -2.348929 | down | PDGFRL |
| A_33_P3219601 | 1.7053704 | up | ABL2 |
| A_33_P3216059 | -2.2134275 | down | ASPN |
| A_21_P0000417 | 1.5931325 | up | SNORD114-20 |
| A_23_P82651 | -1.7671198 | down | NPTX2 |
| A_23_P106844 | 3.3833823 | up | MT2A |
| A_21_P0007907 | 1.8272978 | up | LINC00540 |
| A_23_P77493 | -1.6682904 | down | TUBB3 |
| A_23_P121695 | -12.019378 | down | CXCL13 |
| A_33_P3336038 | 2.1078234 | up | |
| A_33_P3357678 | -2.6267617 | down | LCTL |
| A_23_P302672 | -2.3001962 | down | DDIT4L |
| A_33_P3352970 | 1.9945667 | up | IRAK2 |
| A_24_P59667 | -2.2186131 | down | JAK3 |
| A_23_P107421 | -3.211837 | down | TK1 |
| A_33_P3269924 | 1.7626315 | up | HIP1R |
| A_23_P88404 | -2.2585232 | down | TGFB3 |
| A_23_P53193 | -2.4308362 | down | SYTL2 |
| A_24_P807031 | -1.8642961 | down | ATP6AP1L |
| A_33_P3357658 | 1.8469795 | up | HMGA2 |
| A_23_P417918 | -2.7616024 | down | PENK |
| A_33_P3313283 | -1.9160987 | down | CILP2 |
| A_23_P307310 | -2.5691743 | down | ACAN |
| A_33_P3373765 | -2.209101 | down | DRD4 |
| A_23_P161659 | -2.2976465 | down | SYT13 |
| A_23_P146830 | -2.0724711 | down | SLC25A10 |

| | | | |
|---|---|---|---|
| A_33_P3865368 | -2.8600497 | down | LOC254896 |
| A_33_P3384108 | -1.5629877 | down | SLC19A1 |
| A_23_P253958 | -2.7345374 | down | LRRC17 |
| A_24_P20630 | -1.9027693 | down | LEF1 |
| A_23_P40847 | 1.5296535 | up | CHST2 |
| A_21_P0000241 | -1.5964937 | down | SNORA33 |
| A_33_P3293247 | -1.829082 | down | GPSM1 |
| A_23_P128698 | 1.7282763 | up | SPRY2 |
| A_23_P159907 | -1.5198381 | down | MAGED4B |
| A_23_P64617 | 1.6703097 | up | FZD4 |
| A_24_P854492 | -2.2031581 | down | MIAT |
| A_33_P3258617 | 3.83942 | up | PLIN2 |
| A_23_P374322 | 1.6600087 | up | LACC1 |
| A_23_P347632 | 1.6803648 | up | MTSS1 |
| A_33_P3369844 | -3.3596208 | down | CD24 |
| A_23_P134176 | 1.9052763 | up | SOD2 |
| A_24_P220485 | 1.7158762 | up | OLFML2A |
| A_23_P162918 | -2.1468494 | down | SERPINA3 |
| A_33_P3280721 | 1.6610662 | up | WLS |
| A_32_P154830 | 1.6259212 | up | OSTM1 |
| A_23_P75283 | -2.4628677 | down | RBP4 |
| A_33_P3295358 | 1.5093889 | up | ANGPTL4 |
| A_24_P58337 | 2.2320836 | up | FTH1 |
| A_23_P103775 | -2.0466821 | down | LRRC8C |
| A_33_P3291454 | 2.7580411 | up | CCDC172 |
| A_33_P3225327 | 1.7127436 | up | ATP13A3 |
| A_33_P3368193 | 2.9474494 | up | PNLIPRP3 |
| A_33_P3350488 | -2.383596 | down | NUSAP1 |
| A_23_P37983 | 3.878689 | up | MT1B |
| A_33_P3394517 | 1.7059284 | up | SMG1 |
| A_23_P24129 | 2.3110256 | up | DKK1 |
| A_21_P0000412 | 1.6427149 | up | SNORD114-15 |
| A_24_P10657 | -1.5644559 | down | SLC44A2 |
| A_32_P313405 | 2.6523876 | up | LAMA1 |
| A_23_P74449 | -2.0559578 | down | HPDL |
| A_23_P54840 | 4.0015736 | up | MT1A |
| A_24_P345846 | 1.9497646 | up | ANTXR2 |
| A_32_P202013 | 2.0841515 | up | FAM196A |
| A_23_P34788 | -1.9697758 | down | KIF2C |
| A_23_P94397 | -1.845222 | down | OMD |
| A_23_P139704 | 1.8261888 | up | DUSP6 |
| A_33_P3263274 | 1.5505753 | up | |
| A_33_P3221203 | -2.21771 | down | MMP13 |
| A_23_P59602 | 1.5659834 | up | MIOS |
| A_23_P210210 | 1.7436364 | up | EPAS1 |
| A_22_P00022010 | -2.9775221 | down | FGF11 |
| A_23_P70785 | -1.5065684 | down | AIM1 |
| A_22_P00009049 | -2.738587 | down | LEFTY2 |
| A_32_P234145 | -2.9416072 | down | SHC4 |
| A_22_P00005435 | -3.3791647 | down | lnc-DTYMK-4 |
| A_23_P148410 | 2.022386 | up | FTHL17 |

| | | | |
|---|---|---|---|
| A_23_P365614 | -1.6402571 | down | NOTCH4 |
| A_24_P369232 | -1.597015 | down | CCDC3 |
| A_33_P3318581 | -1.7860152 | down | PLOD2 |
| A_23_P40240 | -1.624729 | down | CTSZ |
| A_23_P258136 | -2.37495 | down | MXRA5 |
| A_23_P134953 | 1.8288531 | up | PLIN2 |
| A_21_P0001959 | 2.0838766 | up | |
| A_23_P214144 | -2.8607438 | down | COL10A1 |
| A_23_P70670 | -1.6710173 | down | CD83 |
| A_33_P3420013 | 1.7038668 | up | |
| A_33_P3252333 | -1.6222762 | down | SNORD3B-1 |
| A_33_P3231156 | 3.066879 | up | |
| A_21_P0000422 | 1.6376499 | up | SNORD114-26 |
| A_33_P3216442 | -2.7643728 | down | COL11A2 |
| A_23_P23839 | -2.1654763 | down | LGR6 |
| A_23_P69179 | -2.0337882 | down | P3H2 |
| A_23_P431853 | 1.834361 | up | ND2 |
| A_33_P3387831 | -2.3296032 | down | CENPM |
| A_23_P95060 | -2.0247326 | down | EPHB3 |
| A_23_P63789 | -1.7675319 | down | ZWINT |
| A_23_P23296 | -2.7838058 | down | PKP1 |
| A_23_P122068 | -2.2981315 | down | C1QTNF3 |
| A_23_P319423 | -3.5883856 | down | KCNK5 |
| A_33_P3272773 | 1.605454 | up | ZNF217 |
| A_23_P153964 | 2.3874896 | up | INHBB |
| A_24_P366082 | 1.6503613 | up | MICAL3 |
| A_32_P191895 | 1.6260769 | up | |
| A_24_P681011 | 2.0677953 | up | HIPK2 |
| A_33_P3639068 | 2.6574302 | up | GFRA1 |
| A_33_P3399788 | -2.6643841 | down | SERPINA3 |
| A_21_P0000425 | 1.5529333 | up | SNORD114-28 |
| A_24_P150791 | -1.5611042 | down | JPH3 |
| A_24_P69095 | -1.5787603 | down | ENC1 |
| A_24_P363408 | -3.002074 | down | HEY2 |
| A_33_P3241269 | 1.9079748 | up | CES1 |
| A_33_P3256894 | 1.6003658 | up | FAM21C |
| A_24_P672240 | -2.327315 | down | FRMPD4 |
| A_33_P3372099 | -1.8462248 | down | DDIT4L |
| A_23_P500464 | -4.7316175 | down | COL2A1 |
| A_33_P3370094 | 1.6148981 | up | MME |
| A_23_P7727 | -3.4444466 | down | HAPLN1 |
| A_21_P0011609 | 2.7657998 | up | LOC100130520 |
| A_22_P00000185 | -1.7356641 | down | |
| A_24_P827037 | -2.0867977 | down | LRRC15 |
| A_23_P9614 | -2.5749652 | down | NDUFA4L2 |
| A_32_P62997 | -2.4002347 | down | PBK |
| A_33_P3223116 | 1.8566751 | up | HIPK2 |
| A_24_P52697 | -2.230038 | down | H19 |
| A_23_P14302 | 1.8794309 | up | LINC00341 |
| A_23_P99386 | -3.6449013 | down | TNFSF11 |
| A_23_P216429 | -2.788618 | down | ASPN |

| | | | |
|---|---|---|---|
| A_32_P169114 | -1.7238501 | down | GRIN2A |
| A_23_P154037 | 1.8554447 | up | AOX1 |
| A_23_P205611 | 1.5667835 | up | GMFB |
| A_23_P114983 | 3.3433528 | up | TRIM63 |
| A_24_P96403 | -1.5832856 | down | RUNX1 |
| A_23_P13382 | -1.7970603 | down | LSP1 |
| A_24_P55295 | -1.8377581 | down | GJA1 |
| A_21_P0000225 | -1.5050081 | down | SNORD83B |
| A_23_P29684 | -1.867384 | down | VILL |
| A_21_P0008856 | 1.6062492 | up | PEAK1 |
| A_22_P00007531 | -1.8545805 | down | LEF1-AS1 |
| A_33_P3255824 | -2.5166824 | down | |
| A_23_P125505 | -3.4709384 | down | PPEF1 |
| A_33_P3419945 | 1.8168601 | up | |
| A_22_P00011907 | -1.6757473 | down | lnc-PIK3R1-5 |
| A_33_P3393971 | -2.4798152 | down | PKP1 |
| A_33_P3329522 | -2.8173382 | down | LRRC17 |
| A_33_P3420466 | -2.5521631 | down | MATN3 |
| A_23_P150343 | -3.5879805 | down | SLN |
| A_23_P25674 | -1.6811776 | down | CKB |
| A_22_P00010174 | -1.5751045 | down | FGD5-AS1 |
| A_33_P3224809 | 2.0092428 | up | IL17RA |
| A_23_P45786 | -2.83676 | down | COL9A2 |
| A_21_P0014651 | -1.8241392 | down | LOC100129203 |
| A_33_P3290707 | 1.9766794 | up | MME |
| A_33_P3280845 | -1.6678222 | down | THY1 |
| A_32_P29615 | 1.8962659 | up | ZNF468 |
| A_24_P62505 | -1.6618257 | down | COLGALT2 |
| A_23_P92441 | -1.8166896 | down | MAD2L1 |
| A_23_P57379 | -2.6289053 | down | CDC45 |
| A_33_P3341787 | 1.5730774 | up | |
| A_23_P19590 | -1.6357003 | down | EZR |
| A_33_P3352407 | 2.1294029 | up | |
| A_21_P0000035 | -1.7244037 | down | TMEM217 |
| A_23_P2920 | -2.156265 | down | SERPINA3 |
| A_23_P38795 | 1.9981962 | up | FPR1 |
| A_33_P3404052 | 1.8713673 | up | TNFAIP8 |
| A_23_P10206 | -2.1815712 | down | HAS2 |
| A_21_P0000402 | 1.7778257 | up | SNORD114-5 |
| A_24_P119609 | 1.6217246 | up | MYO1D |
| A_23_P256047 | -2.3039114 | down | ANKEF1 |
| A_23_P58082 | -1.6145083 | down | CCDC80 |
| A_23_P167159 | -2.2205184 | down | SCRG1 |
| A_23_P371239 | -2.1975982 | down | CMIP |
| A_23_P152235 | 1.6088113 | up | IRX3 |
| A_33_P3278033 | -1.9780755 | down | TUSC1 |
| A_23_P137381 | -1.7172724 | down | ID3 |
| A_23_P401 | -1.6074336 | down | CENPF |
| A_23_P502464 | -4.0967383 | down | NOS2 |
