## Supplementary material for "Epigenetic Regulation of Chondrogenesis: JMJD3 and UTX as Key Targets for Gene-Modified Mesenchymal Stem Cell Therapy in Cartilage Tissue Engineering": suppl table 2

**Suppl table 2 : Gene Ontology (GO) biological process enrichment**

| #Term ID | Term description | Observed gene count | Background gene count | Strength | Signal | False discovery rate | Matching proteins in your network (labels) |
|---|---|---|---|---|---|---|---|
| GO:0030198 | Extracellular matrix organization | 32 | 278 | 0.82 | 1.74 | 6.35E-12 | CCDC80, FERMT1, IBSP, SOX9, MATN2, SULF1, MMP13, LUM, ADAMTS16, IHH, PTX3, HAS2, COL10A1, ADAMTSL2, SMOC2, FMOD, COL27A1, COLGALT2, COL9A2, OLFML2A, COL11A2, COL5A2, COL2A1, ADAMTSL1, LOXL2, SLC39A8, MATN3, ACAN, ADAMTS9, MIA, COL9A3, GREM1 |
| GO:0001503 | Ossification | 26 | 287 | 0.71 | 1.22 | 5.86E-08 | SFRP1, IBSP, IGFBP5, TGFB3, SOX9, MMP13, LEF1, IHH, RUNX1, WNT11, FGFR3, LRRC17, FZD9, COL11A2, COL5A2, ASPN, COL2A1, IGF1, SPP1, TNFSF11, PENK, PTH1R, PHOSPHO1, SP7, DLX5, CLEC3A |
| GO:0001501 | Skeletal system development | 34 | 513 | 0.58 | 1.04 | 1.35E-07 | SFRP1, MDFI, TGFB3, SOX9, SULF1, MMP13, HAPLN1, GJA1, FRZB, IHH, RUNX1, HAS2, WNT11, COL10A1, HOXD1, FGFR3, LRRC17, COL27A1, ETS2, RBP4, COL9A2, COL11A2, COL5A2, OGN, COL2A1, IGF1, TNFSF11, MATN3, ACAN, PTH1R, PHOSPHO1, HMGA2, DLX5, CLEC3A |
| GO:0010273 | Detoxification of copper ion | 6 | 14 | 1.39 | 0.95 | 0.00021 | MT2A, MT1A, MT1E, MT1B, MT1X, MT1HL1 |
| GO:0009888 | Tissue development | 74 | 1723 | 0.39 | 0.91 | 2.59E-09 | KRT14, CTSZ, SFRP1, IBSP, IGFBP5, SOX9, KRT12, SULF1, MMP13, NOX4, EPAS1, CSTA, LEF1, SEMA3A, CPT1A, ADAMTS16, GJA1, PPARG, FRZB, IHH, RUNX1, HAS2, TUBB3, MSC, MTSS1, WNT11, IRX3, DACT1, EZR, FGFR3, LRRC17, ATF3, ADAMTSL2, WLS, SORBS2, COL27A1, ETS2, CES1, DMD, HEY2, RBP4, MFSD2A, DKK1, ID3, COL11A2, NOTCH4, ASPN, OGN, SPRY2, COL2A1, AKR1C1, LOXL2, LAMA1, CRYGS, IGF1, FOLR1, SPP1, CDK1, TNFSF11, MATN3, ACAN, PTH1R, PHOSPHO1, ADAMTS9, MEGF10, CEBPA, HMGA2, SP7, ITGB3, MYC, CD24, KITLG, DLX5, GREM1 |
| GO:0051093 | Negative regulation of developmental process | 47 | 933 | 0.46 | 0.9 | 2.07E-07 | CHADL, NFKBIA, FERMT1, SFRP1, SPARC, IGFBP5, SOX9, SULF1, LEF1, SEMA3A, PLK2, THY1, PPARG, FRZB, IHH, RUNX1, WNT11, IRX3, CXCL14, FGFR3, TP53INP1, LRRC17, DNM3, LDLRAD4, DMD, ASPM, HEY2, RBP4, DKK1, ID3, COL5A2, NOTCH4, ASPN, ID1, SPRY2, RGCC, LOXL2, IGF1, SPP1, NFATC1, ADAMTS9, CEBPA, JAK3, SOD2, ITGB3, MYC, GREM1 |
| GO:2000026 | Regulation of multicellular organismal development | 62 | 1389 | 0.41 | 0.88 | 4.29E-08 | CHADL, NFKBIA, FERMT1, SFRP1, CTSC, SPARC, TGFB3, SOX9, SULF1, LEF1, SEMA3A, PLK2, NTRK2, DUSP6, THY1, CXCL13, PPARG, S100A1, FRZB, IHH, RUNX1, ANGPTL4, TENT5A, WNT11, EPHB3, ZMIZ1, LRRC17, FZD9, SMOC2, ASPM, HEY2, RBP4, DKK1, NOTCH4, ASPN, OMD, ID1, SPRY2, CD83, RGCC, LOXL2, LAMA1, IGF1, SPP1, CDK1, TNFSF11, HIPK2, ATP2B1, PHOSPHO1, MME, ADAMTS9, CEBPA, JAK3, HMGA2, ITGB3, MYC, TNFAIP3, CD24, NR2C2, SPAAR, KITLG, GREM1 |
| GO:0030111 | Regulation of Wnt signaling pathway | 23 | 326 | 0.61 | 0.85 | 1.80E-05 | FERMT1, SFRP1, MDFI, SOX9, SULF1, LEF1, FRZB, NKD2, GPRC5B, WNT11, DACT1, FZD9, WLS, LGR6, ASPM, DKK1, FOLR1, NFATC1, FZD4, TLE2, TNFAIP3, DLX5, GREM1 |
| GO:0046688 | Response to copper ion | 8 | 40 | 1.06 | 0.84 | 0.00033 | MT2A, MT1A, MT1E, MT1B, LOXL2, MT1X, CDK1, MT1HL1 |
| GO:0002062 | Chondrocyte differentiation | 11 | 85 | 0.87 | 0.83 | 0.00021 | SOX9, SULF1, IHH, RUNX1, FGFR3, COL27A1, COL11A2, COL2A1, ACAN, PTH1R, HMGA2 |
