## Supplementary material for "Epigenetic Regulation of Chondrogenesis: JMJD3 and UTX as Key Targets for Gene-Modified Mesenchymal Stem Cell Therapy in Cartilage Tissue Engineering": suppl table 3

**Suppl table 3: KEGG enrichment**

| #Term ID | Term description | Observed gene count | Background gene count | Strength | Signal | False discovery rate | Matching proteins in your network (labels) |
|---|---|---|---|---|---|---|---|
| hsa04550 | Signaling pathways regulating pluripotency of stem cells | 13 | 141 | 0.72 | 0.67 | 0.00089 | INHBB, WNT11, HOXD1, FGFR3, FZD9, LEFTY2, ID3, ID1, IGF1, JAK3, FZD4, MYC, DLX5 |
| hsa04978 | Mineral absorption | 8 | 57 | 0.91 | 0.63 | 0.0025 | MT2A, FTH1, MT1A, MT1B, MT1X, SLC11A2, ATP2B1, MT1HL1 |
| hsa05200 | Pathways in cancer | 22 | 515 | 0.39 | 0.36 | 0.0182 | NFKBIA, TGFB3, EPAS1, LEF1, PPARG, RUNX1, WNT11, NOS2, GNG2, FGFR3, FZD9, STAT4, HEY2, NOTCH4, LAMA1, IGF1, CEBPA, JAK3, FZD4, PLEKHG5, MYC, KITLG |
| hsa04350 | TGF-beta signaling pathway | 8 | 91 | 0.7 | 0.4 | 0.0208 | TGFB3, INHBB, FMOD, LEFTY2, ID3, ID1, MYC, GREM1 |
| hsa04512 | ECM-receptor interaction | 8 | 88 | 0.72 | 0.4 | 0.0208 | IBSP, COL9A2, COL2A1, LAMA1, HMMR, SPP1, ITGB3, COL9A3 |
| hsa05224 | Breast cancer | 10 | 146 | 0.59 | 0.39 | 0.0209 | LEF1, WNT11, SHC4, FZD9, HEY2, NOTCH4, IGF1, TNFSF11, FZD4, MYC |
| hsa04310 | Wnt signaling pathway | 10 | 154 | 0.57 | 0.36 | 0.0265 | SFRP1, LEF1, NKD2, WNT11, FZD9, LGR6, DKK1, NFATC1, FZD4, MYC |
| hsa04974 | Protein digestion and absorption | 8 | 100 | 0.66 | 0.38 | 0.0265 | COL10A1, COL27A1, KCNK5, COL9A2, COL5A2, COL2A1, MME, COL9A3 |
